# Laminar compartmentalization of attention modulation in area V4 aligns with the demands of visual processing hierarchy in the cortex

**DOI:** 10.1101/2021.02.15.431312

**Authors:** Xiang Wang, Anirvan S. Nandy, Monika P. Jadi

## Abstract

Contrast is a key feature of the visual scene that aids object recognition. Attention has been shown to selectively enhance the responses to low contrast stimuli in visual area V4, a critical hub that sends projections both up and down the visual hierarchy. Veridical encoding of contrast information is a key computation in early visual areas, while later stages encode higher level features that benefit from improved sensitivity to low contrast. How area V4 meets these distinct information processing demands in the attentive state is not known. We found that attentional modulation of contrast responses in area V4 is cortical layer and cell-class specific. Putative excitatory neurons in the superficial output layers that project to higher areas show enhanced boosting of low contrast information. On the other hand, putative excitatory neurons of deep output layers that project to early visual areas exhibit contrast-independent scaling. Computational modeling revealed that such layer-wise differences may result from variations in spatial integration extent of inhibitory neurons. These findings reveal that the nature of interactions between attention and contrast in V4 is highly compartmentalized, in alignment with the demands of the visual processing hierarchy.

## INTRODUCTION

Voluntary attention is essential for sensory guided behavior and memory formation (Petersen and Posner, 2012). Failures in sensory processing and selective attention are aspects of many mental illnesses, including schizophrenia and mood disorders (Fioravanti et al., 2005; McIntyre et al., 2010; Neuchterlein et al., 1991). Visual spatial attention plays a critical role in visual sensory processing: It allows improved perception of behaviorally relevant target stimuli among competing distractors by boosting the apparent visibility of the target (Carrasco et al., 2004). At the neuronal level, attention modulates the activity of cortical neurons that encode an attended visual stimulus at various stages of visual processing (Bisley and Goldberg, 2003; Ghose and Maunsell, 2008; Moran and Desimone, 1985; Motter, 1993; Reynolds et al., 1999; Treue and Martinez Trujillo, 1999; Treue and Maunsell, 1996). In visual areas such as V4 and MT, attention modulates neuronal mean firing rates, increases their firing reliability, and reduces the co-variability among pairs of neurons (Cohen and Maunsell, 2009; Mitchell et al., 2007, 2009; Reynolds and Chelazzi, 2004; Treue and Martinez Trujillo, 1999). However, the computational principles that underlie the activity of neuronal populations that represent both sensory information and the attentional state remain poorly understood (Moore and Zirnsak, 2017; Reynolds and Chelazzi, 2004).

Object recognition is mediated by a hierarchy of cortical visual processing areas that form the ventral visual stream. Contrast is a key feature of the visual scene that aids object recognition, and the encoding of contrast information is one of the most important computations performed by early visual areas. On the other hand, visual features represented in higher areas such as the inferotemporal (IT) cortex benefit from improved sensitivity to low contrast stimuli (Avidan et al., 2002; Rolls and Baylis, 1986). Visual area V4 is a critical hub in the ventral stream that sends feedforward projections to areas such as IT and feedback projections to early visual processing areas (Anderson and Martin, 2006; Douglas and Martin, 1991; Van Essen and Maunsell, 1983). Attention has been shown to selectively enhance the responses to low contrast stimuli (Martinez-Trujillo and Treue, 2002; Reynolds et al., 2000). Attention mediated selective enhancement of low contrast features is thought to aid invariant representations in higher object recognition areas downstream of V4 (Roe et al., 2012). However, such a bias in the attention-modulated feedback from V4 to upstream visual areas can disrupt the contrast-based feature extraction functions of these stages. How area V4 meets these distinct information processing demands of the visual processing hierarchy is not known. While attention can enhance V4 responses in a contrast-independent manner (response gain) under certain experimental conditions (Williford and Maunsell, 2006), an understanding of robust mechanisms of feedback from V4 that does not interfere with the contrast landscape of scene representations in early visual areas remains elusive.

One possibility is that distinct subpopulations in V4 mediate these functional demands. Indeed, the sensory cortical sheet, including area V4, is not a homogeneous piece of tissue along its depth; rather, it has a six-layered or laminar structure made up of multiple cell classes, of both excitatory and local inhibitory kind, with largely stereotypical anatomical connectivity between and within layers (Douglas and Martin, 2004). Layer 4 (the *input* layer) is the primary target of projections carrying visual information from early areas, such as V1, V2, and V3 (Felleman and Van Essen, 1991; Ungerleider et al., 2008). Visual information is then processed by local neural subpopulations as it is sent to layers 2/3 (the *superficial* layer) and layers 5/6 (the *deep* layer), which serve as output nodes in the laminar circuit (Hirsch and Martinez, 2006; Rockland and Pandya, 1979). The superficial layers feed information forward to downstream visual areas, such as IT (Borra et al., 2010; Distler et al., 1993), whereas the deep layers send feedback information to upstream early visual areas (Callaway, 1998; Gattass et al., 2014; Mehta et al., 2000; Ungerleider et al., 2008). This anatomical organization suggests distinct functional roles (D’Souza and Burkhalter, 2017), and differential attentional modulation of sensory representation among cell-class and layers-specific neural subpopulations. In support of this idea, a recent study of simultaneous depth recordings in visual area V4 has shown layer-specific attentional modulation of average neuronal responses, reliability of responses, and correlations between responses of pairs of neurons (Nandy et al., 2017). Therefore, to fully understand the attentional modulation of sensory computations, it is essential to investigate the modulation of sensory representation in these subpopulations. Our broad hypothesis is that the attentional modulation of contrast computations in area V4 is not homogeneous, but rather is layer- and cell-class specific and that these differences reflect the different computational demands on these subpopulations. Considering their key contribution to feedback projections to early visual areas, we specifically expect that projection neurons in the deep layers show uniform attentional modulation across all contrasts in order to minimally impact the faithful representation of contrast landscape in their target areas.

In this study, we characterized layer- and cell-class specific neural subpopulations from extracellular simultaneous laminar recordings of single neurons within area V4 of macaque monkeys performing an attention-demanding task. Using unsupervised clustering techniques on spiking properties, we distinguished five functional clusters of neurons. We distinguished layer identities – superficial, input or deep – of these neurons using features of local field potentials. To test our hypothesis, we characterized the attentional modulation of contrast response functions in these sub-populations. We interpreted our findings within a computational framework of attentional modulation of contrast responses (Reynolds and Heeger, 2009), which yielded predictions for distinct mechanistic roles of these neural subpopulations in attentive perception.

## RESULTS

In the primate visual system, cortical sensitivity to features such as luminance contrast varies with the locus of spatial attention; contrast response functions (CRF) of cortical neurons are measured to quantify this dependence (Kastner and Ungerleider, 2000; Reynolds and Chelazzi, 2004; Reynolds et al., 2000). However, the laminar- and cell-class specific dependence of CRF on the attentive state is not known. Using linear array electrodes, we recorded neuronal activity from well-isolated single units, multi-unit clusters, and local field potentials (LFPs) in visual area V4 of two rhesus macaques (right hemisphere in monkey A, left hemisphere in monkey C) during an attention demanding orientation change detection task (Figure 1A, B; see Methods). We used current source density (CSD) analysis to identify different laminar compartments (*superficial*, *input*, and *deep*), and assigned isolated single units to one of the three layers (see Methods). In the main experiment, we presented a sequence of paired Gabor stimuli with different contrasts (Figure 1B); one stimulus was presented inside the receptive fields (RFs) of the recorded neurons and the other at an equally eccentric location across the vertical meridian. Attention was cued either to the stimuli within the neurons’ RFs (“attend-in”) or to the stimuli in the contralateral visual hemifield (“attend-away”).

**Figure 1.**
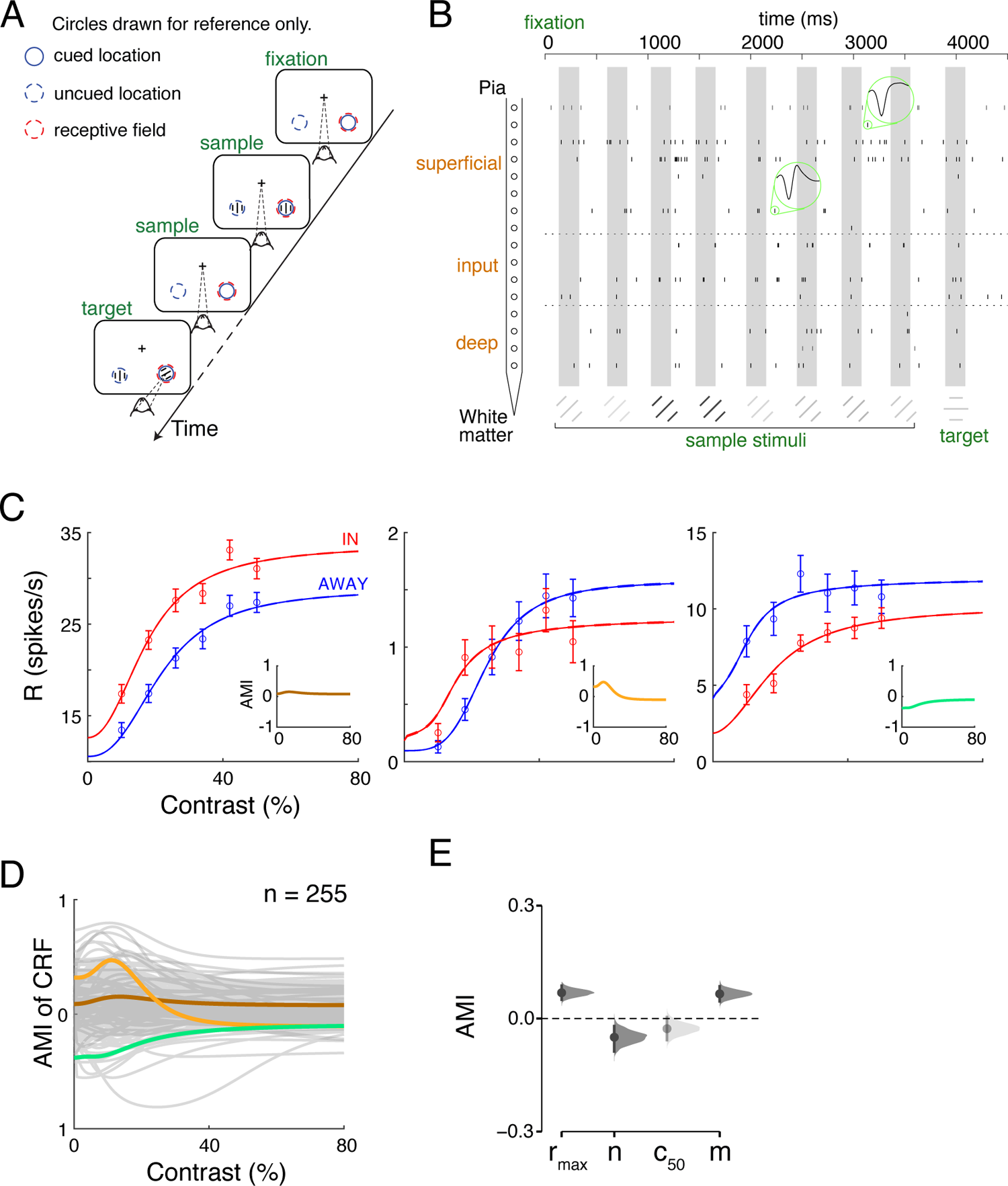
Attentional modulation of Contrast Response Function. (A) Orientation change detection task. While the monkey maintained fixation, two oriented Gabor stimuli were flashed on and off at two locations: one within the RF overlap region of the recorded V4 column and the other at a location of equal eccentricity across the vertical meridian. The covert attention of the monkey was cued to one of the two locations. One of the two stimuli changed its orientation at an unpredictable time. The monkey was rewarded for making a saccade to the location of orientation change (95% probability of change at the cued location; 5% probability at uncued location [foil trials]). If no change happened (catch trials), the monkey was rewarded for maintaining fixation. (B) An example trial showing the single-unit signals in the attend-in condition. The time axis is referenced to the appearance of the fixation spot. Spikes (vertical ticks) in each channel come from the single unit with the highest spike rate in this trial. The grey boxes depict stimulus presentation epochs. In this particular trial, 8 sample stimuli with different contrasts were presented, followed by a target stimulus flash with an orientation change that the monkey responded to correctly. Two different waveforms were shown for two single units. (C) The best-fitting contrast response functions of three example neurons in “attend in” and “attend away” conditions. Mean ± SEM. Insets show the attentional modulation indices calculated as a function of contrast. (D) The AMI as a function of contrast for each of the 255 visually responsive single units, with the three example units in (C) highlighted. (E) The Cumming estimation plot shows the bootstrap sampling distribution of mean AMI for each parameter. The average is depicted as a dot. Each 95% confidence interval (CI) is indicated by the ends of the vertical error bars. The faded color represents that the 95% CI include 0.

### Attentional Modulation of Contrast Response Function

To examine the effects of attention on individual neurons, we used the method of ordinary least squares to fit each neuron’s contrast responses from both attentional states to a hyperbolic ratio function (Figure 1C). This function is described by four parameters: *r_max_*, *C_50_*, *m*, and *n*, where *r_max_* is the attainable maximum response, *C_50_* is the contrast at which neuronal response is half-maximal, *m* is the baseline activity, and *n* describes the nonlinearity of the function. Attention effects differ considerably for individual neurons. Attention either enhances or suppresses neuronal responses at different contrast levels (Figure 1D). We quantified the effect of attention on every recorded neuron by computing the attentional modulation index (AMI) using contrast responses from both attention conditions (see Methods). We saw a significant variance of AMI values at each contrast level (Figure 1E). We also examined how attention impacts the values of best-fitting parameters (Figure 1F). The mean AMIs for *r_max_* and *m* are significantly higher than zero (Mann-Whitney U test, p < 0.01 for both distributions), which is consistent with previous observations in V4 (Williford and Maunsell, 2006). The same percentage change in *r_max_* and *m* (15% increase) supports an effect of contrast independent scaling by attention. The average modulations of *C_50_* and *n* are significantly smaller than zero (Mann-Whitney U test, p < 0.01 for *C_50_* and p ≪ 0.01 for *n*), suggesting an increased sensitivity to low contrast stimuli and a reduction in the sensitivity to contrast change, respectively. The bootstrap sampling distributions of the mean difference from 0 support the average attention effects on *r_max_*, *n* and *m* (Figure 1G). These results indicate that the overall effect of attention on V4 neuron responses cannot be simply explained as selective boosting of low contrast. It is a combination of modulations in multiple parameters of the contrast response function (Figure 1F, G).

### Classification of Single Units Using Electrophysiological Features

To investigate whether attention modulates different classes of neurons uniformly or differentially, we characterized classes of single units based on the peak-to-trough duration (PTD). Properties of the action potential waveform, especially the PTD, have been extensively used to classify neurons into narrow-(putative inhibitory) and broad-spiking (putative excitatory) cells (Constantinidis and Goldman-Rakic, 2002; Diester and Nieder, 2008; Hussar and Pasternak, 2009; Johnston et al., 2009; Kaufman et al., 2010; Mitchell et al., 2007; Wilson et al., 1994). The shapes of average spiking waveform for all single units in our data were also highly variable (Figure 2B). We exploited the information structure in the entire waveforms by applying principal component analysis (PCA). The correlation pattern between the first two components of the PCA (cumulative percentages of explained variance: 59.62%, 83.10%) supported the idea that neurons can be separated into meaningful clusters by waveform shape measures (Figure 2C).

**Figure 2.**
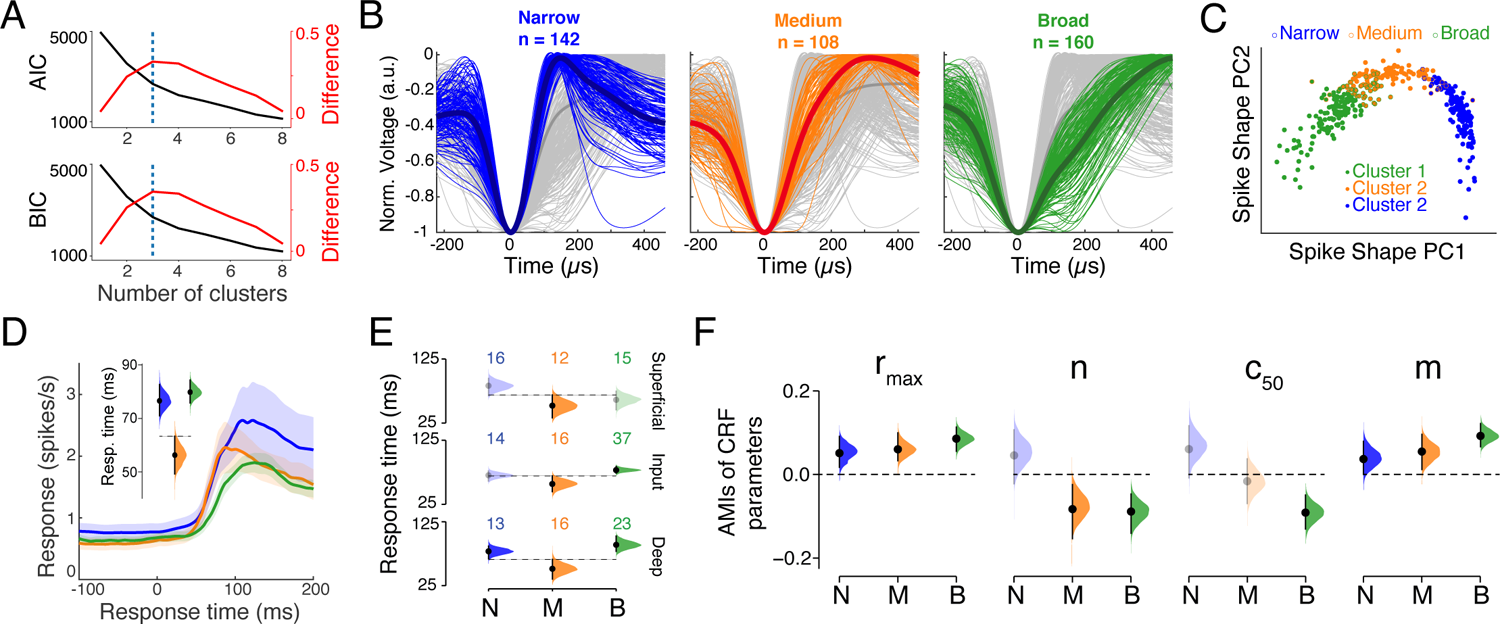
Classification of Single Units Using Waveform Widths. (A) AIC and BIC for different number of clusters. The elbow point (purple) was detected by the kneedle algorithm as the local maxima of the difference curve (red) computed from the normalized AIC or BIC values. (B) Mean waveforms for all 410 single units colored by their cluster identies. Waveforms were smoothed using spline interpolation and their heights were normalized to help compare spike widths. The mean waveform averaged across all units was shown in dark grey and the mean waveforms for each cluster were shown in dark colors. (C) In the principal component space of spike shape, single units were colored either based on their spike width classifications (open circles; Narrow, Medium, Broad) or by running the *k*-means clustering algorithm with the first 2 PCs (closed circles). The clusters generated from the peak-to-trough duration match the ones classified by spike shape PCs, suggesting that peak-to-trough duration is an efficient measure to capture the variance of spike shape data. (D) Lines and shading show mean ± SEM firing rates for 3 classes around the stimulus onset, averaged across visually responsive neurons (Narrow, *n* = 56; Medium, *n* = 64; Broad, *n* = 87 neurons). The inset shows the bootstrap sampling distribution of the mean response time for each cluster. Response time is the interval between the stimulus onset and the time when the post-stimulus firing rate is greater than the 95% CI of the pre-stimulus mean firing rate. The dashed line indicates the upper bound of the 95% CI of Medium neurons. (E) Layer-wise distributions of mean response time for 3 classes (Narrow, N; Medium, M; Broad, B). The number of units is on top of each distribution. Dashed lines indicate the upper bounds of 95% CIs of Medium neurons. Distributions with CIs overlapping with the Medium class are shown in faded colors. (F) Bootstrap sampling distributions of AMIs of CRF parameters for each cell class. Distributions with CIs including 0 are displayed in faded colors. The CRF parameters were only available for visually responsive single units.

We used a meta-clustering analysis based on the *k*-means clustering algorithm (see Methods) to identify clusters of isolated single units (see Methods; Ardid et al., 2015; Hartigan and Wong, 1979). To select the optimal number of clusters (*k*), we computed Akaike information criterion (AIC) and Bayesian information criterion (BIC) for each *k*. Elbow points of two criteria (see Methods) supported the selection of the 3-cluster solution (Figure 2A). Narrow-spiking cells become a cluster by themselves, while a medium-spiking cluster emerged from those previously classified as broad-spiking cells (Mitchell et al., 2007; Nandy et al., 2017; Figure 2B).

One of the assumptions we made to use the PTD as a clustering feature was that it captures a significant amount of the variations of neurons’ spiking waveforms. We tested this assumption by clustering neurons in the principal component space of the AP waveform and comparing them with neuronal groups classified by their PTD. We found that the three clusters separated by waveform principal components were consistent with those classified by the spike width (Figure 2C).

The clusters differ in terms of their firing rates (Supp. Figure 2A). Although not statistically significant, narrow-spiking neurons exhibited higher firing rates than medium- and broad-spiking clusters when averaged across layers (mean 10.1 Hz compared to 8.9 Hz and 6.5 Hz; Mann-Whitney U test, p_narrowómedium_ = 0.44, p_narrowóbroad_ = 0.058). It is in agreement with previous findings that narrow-spiking neurons, considered putative inhibitory interneurons, show higher firing rates than broad-spiking neurons, thought to be putative excitatory pyramidal cells (Connors and Gutnick, 1990; McCormick et al., 1985; Mitchell et al., 2007; Nowak et al., 2003; Povysheva et al., 2006).

We also performed a timing analysis to examine the response time of neuronal classes to visual stimuli (see Methods). For each single unit, we defined the response time as its post-stimulus delay at which the firing rate surpassed the upper bound of the 95% confidence interval (CI) calculated from its pre-stimulus spontaneous activity. We then averaged the response time across units within a cluster (Figure 2D) and within a layer (Figure 2E). When averaged across layers, medium-spiking cells showed shorter response time than the other two classes (Figure 2D). Especially, medium-spiking neurons exhibited faster response than broad-spiking cells in the input and deep layers (Figure 2E). These results further support that medium-spiking neurons represent a distinct subpopulation from broad-spiking cells, and the faster response provides strong evidence that the medium-spiking subpopulation might be the first group of cells that receive visual information in the input and deep layers.

### Cell-Class and Layer-Specific Attentional Modulation

We next examined how attention modulates contrast responses for each cell class. We first computed the AMIs of best-fitting CRF parameters for every cell class. The pattern of modulations of CRF parameters was distinct for individual cell classes (Figure 2F). All three clusters showed significant positive modulations of *r_max_* and *m*. Medium and Broad clusters are negatively modulated in their nonlinearity parameter *n*, an effect that enhances low-contrast responses and suppresses high-contrast responses (reduced overall contrast sensitivity). Notably, only the broad-spiking neurons exhibited a change of *c_50_* by attention, implying a selective enhancement of responses to low contrast stimuli. This effect was novel to the Broad cluster and not revealed in the analysis of unclassified neurons (Figure 1G). None of the other two cell classes – Narrow and Medium – showed a significant modulation of *c_50_* by attention, an effect that matched the analysis of unclassified neurons (Figure 1G).

To further investigate the cell-class specific attentional modulation at each contrast level, we computed the AMI as a function of contrast using CRFs from both attentional states for every single unit and averaged AMIs across single units within a cluster (Figure 3A, left panel). The attentional modulation of *c_50_* and *n* (Figure 2F) suggests that medium- and broad-spiking neurons are more strongly modulated by attention in the range of low contrast than the range of high contrast. Indeed, we found that the AMIs of Medium and Broad clusters were relatively dependent on contrast, whereas the Narrow cluster appeared to be modulated by attention in a contrast-independent manner (Figure 3A, left panel). When averaged across contrasts, attention positively modulated firing rates for all cell classes (Figure 3A, right panel). Broad-spiking cells showed stronger attentional modulation than medium- and narrow-spiking cells. To quantify the contrast dependence of attentional modulation for each single unit, we first averaged the AMIs within the low-contrast and the high-contrast ranges with the threshold set at each unit’s best-fitting *c_50_* parameter. We then defined the contrast dependence index (CDI) of a single unit as the difference between the two average AMIs normalized by the AMI averaged across all contrasts (see Methods). Contrast-independent modulation would then result in CDI = 0, reflecting a pure scaling effect of attention on the CRF. A positive CDI would indicate a more robust attentional modulation at the low-contrast range. A negative CDI would suggest a stronger attention effect on neural responses at the high-contrast range (Figure 3B). We examined the CDI distribution within each cell class and found that the mean CDI of the Narrow class was around zero. However, Medium and Broad clusters exhibited more positive CDIs (Figure 3C). These results are consistent with our findings of AMIs of CRF parameters for each cell class (Figure 2F), confirming that attention modulated narrow-spiking neuron responses regardless of the stimulus contrast. On the other hand, the attention effects on Medium, and Broad clusters were dependent on contrast and were more robust in the low-contrast range.

**Figure 3.**
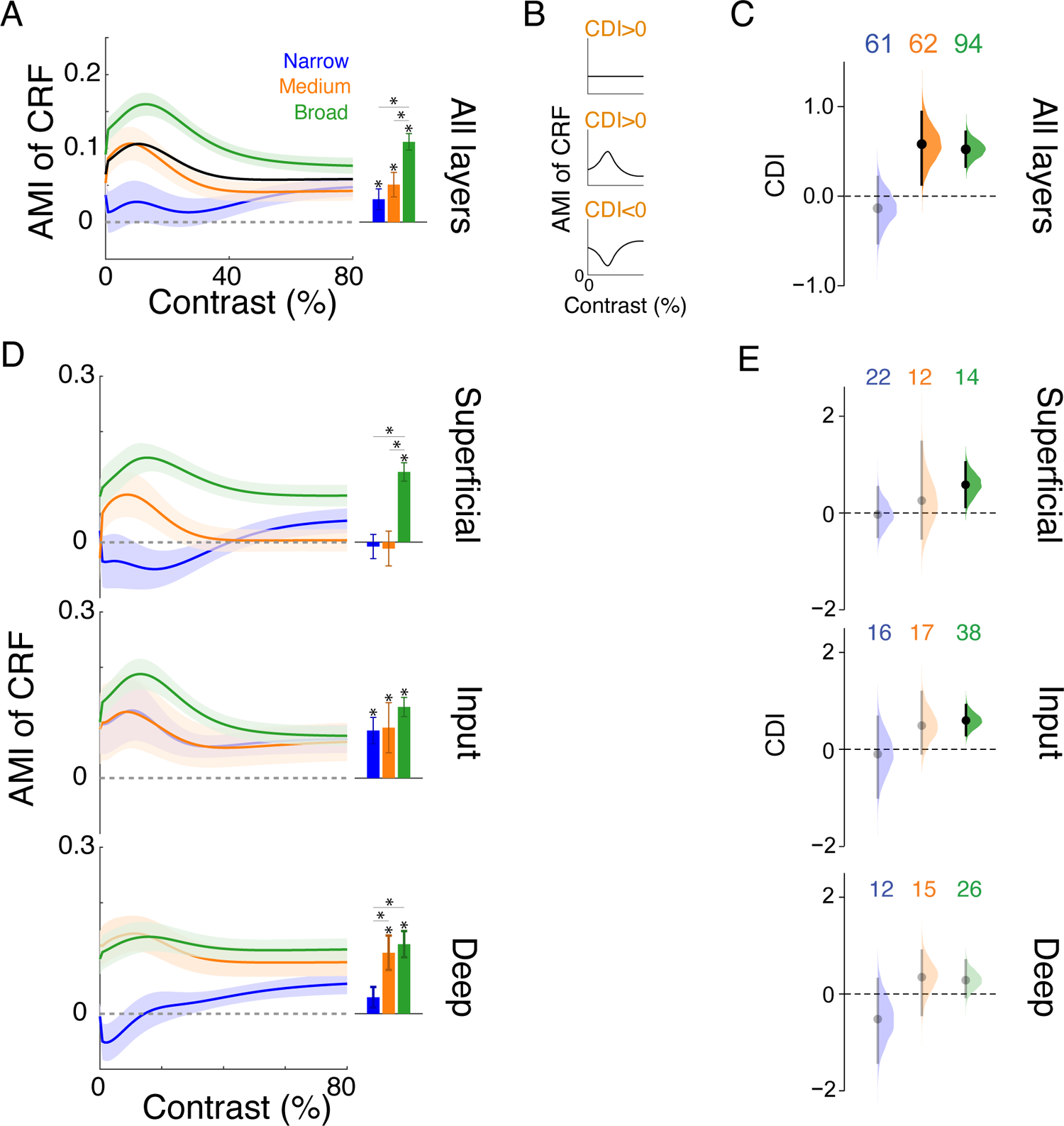
Contrast Dependency of AMI is Cell-Class and Layer-Specific. (A) Left: the AMI of contrast responses as a function of contrast averaged across visually responsive single units in each cluster. Mean ± SEM. The population mean is shown in black. Right: the mean AMI averaged across contrasts for each cluster. Asterisk indicates either the distribution is significantly different from zero or two distributions are significantly different (Mann-Whitney U test, p < 0.05). (B) To quantify the contrast dependence of attentional modulation, we averaged the AMI for a single unit within the low-contrast range or the high-contrast range (using the *c_50_* as the low- to high-contrast threshold). We then defined the contrast dependence index (CDI) as the difference between the average AMI within the low-contrast range and that within the high-contrast range, normalized by the mean AMI across the whole contrast range. The schematic shows the interpretation of different ranges of CDI in terms of the AMI. (C) The bootstrap sampling distribution of the mean CDI for each cell class combined across layers. The number of units excluding outliers for each cell class is shown on the top. Distributions with CIs inclusive of 0 are shown in faded colors. (D) Layer-wise AMI (mean ± SEM) of contrast responses for each cell class as a function of contrast (left) or averaged across contrasts (right). Asterisk indicates either the distribution is significantly different from zero or two distributions are significantly different (Mann-Whitney U test, p < 0.05). (E) Layer-wise bootstrap sampling distribution of the mean CDI for each cell class. Distributions with CIs inclusive of 0 are illustrated in faded colors. The number of units excluding outliers is shown on the top. For the raw data of the layer-wise CDIs, see Supp. Figure 3C.

We further inspected the laminar profile of the attention effect and its contrast dependence for every cell class (Figure 3D, E). When averaged across contrasts, (Figure 3D, right panels), narrow-spiking neurons showed significant attentional modulation only in the input layer, but not in the superficial or deep layer (Figure 3D, right panels, Mann-Whitney U test, p_superficial_ = 0.79, p_input_ < 0.01, p_deep_ = 0.06). On the other hand, the Broad cluster was robustly modulated by attention across all cortical layers (Figure 3D, right panels, Mann-Whitney U test, p_superficial_ ≪ 0.01, p_input_ ≪ 0.01, p_deep_ ≪ 0.01). The AMI difference between these two cell classes is in agreement with the differences between narrow- and “broad”-spiking cells previously reported in these cortical layer (Nandy et al., 2017); it is important to note that the AMI pattern of the Medium cluster across layers was different from the Broad cluster in the superficial layer, again implying that they represent distinct subpopulations (Figure 3D). Two key laminar patterns of contrast dependence emerged from these three clusters. First, the attentional modulation of the Narrow and Medium clusters was independent of contrast across all cortical layers. Second, the Broad cluster exhibited strong contrast dependence and, specifically, significant modulation in the low-contrast range in the superficial and input layers; but its dependence on contrast was not significant in the deep layer (Figure 3E). Also notably, the laminar differences did not emerge when all units in a layer were analyzed as either a single class or more conventionally as narrow vs. “broad” classes (Supp. Figure 3C).

### Laminar network mechanisms of contrast dependence of AMI across layers

We next used computational modeling to gain insights into the possible neural mechanisms underlying the layer- and cell-class specific AMI dependency on stimulus contrast. Variation in CDI across experimental paradigms has been previous observed (Martinez-Trujillo and Treue, 2002; Reynolds et al., 2000; Williford and Maunsell, 2006), and explained by paradigm-specific normalization due to attention (Reynolds and Heeger, 2009). We hypothesized that normalization mechanisms can also explain the layer-specific differences in CDI in our empirical findings (Figure 3D, E). To test this, we first interpreted our results in the context of the normalization model of attention (Reynolds and Heeger, 2009) to generate predictions about layer-specific cortical connectivity that might underlie the variations in CDI. The normalization model of attention proposes a computational principle that accounts for various attention effects on neurons’ contrast response functions (Reynolds and Heeger, 2009). Normalization model assumes that the relative sizes of excitatory receptive field and suppressive field of neurons, and the ‘attention field’ of the experimental paradigm shape the net suppressive drive to individual neurons. The suppressive drive ultimately determines the CDI of individual neurons in a population. We thus investigated the consequences of varying the relative sizes of excitatory receptive field and suppressive field of individual neurons on attentional modulations of CRFs (see Methods). This inquiry was motivated by the observation that neuronal receptive field sizes change along the cortical depth in sensory areas (Gilbert, 1977; Sur et al., 1985; Vaiceliunaite et al., 2013), and based on the assumption that ‘attention field’ sizes are constant for an experimental paradigm.

We simulated the normalization model with different sizes of excitatory receptive field and suppressive field of neurons, and generated neuronal responses to different stimulus contrasts in “attend in” and “attend away” conditions (Figure 4A, top panel). We computed the AMI and the CDI for each combination of size parameters (see Methods). We find that the CDI depends both on the excitatory receptive field size and on the suppressive field size. Holding the attention field size and the stimulus size fixed, a smaller suppressive field or a smaller excitatory receptive field leads to a greater CDI of the attentional modulation (Figure 4A, middle panel). On the other hand, a larger suppressive field or a larger excitatory receptive field results in a smaller CDI (Figure 4A, middle panel). These results hold for a wide range of values of the stimulus size and the attention field size. The pattern is robust when the attention field and the stimulus are both small or large (Supp. Figure 4B, i). The results are also stable for both a linear and saturating transfer function assumption between the stimulus contrast and excitatory drive in the normalization model (Supp. Figure 4B, ii). We also computed the AMI of suppressive drive of neurons for each combination of size parameters. The CDI of model neurons is roughly proportional to the AMI of suppressive drive (Figure 4A, bottom panel). Greater the AMI of suppressive drive, stronger is the CDI of model neurons, and vice versa. Since broad-spiking neurons are putative excitatory pyramidal cells, these results suggest two possible neural mechanisms that explain the laminar profile of CDIs of broad-spiking neurons: the suppressive field size increases along the depth of V4 (Figure 4A, middle panel) or the excitatory receptive field is more extensive in the deeper layer of V4 (Supp. Figure 4C).

**Figure 4.**
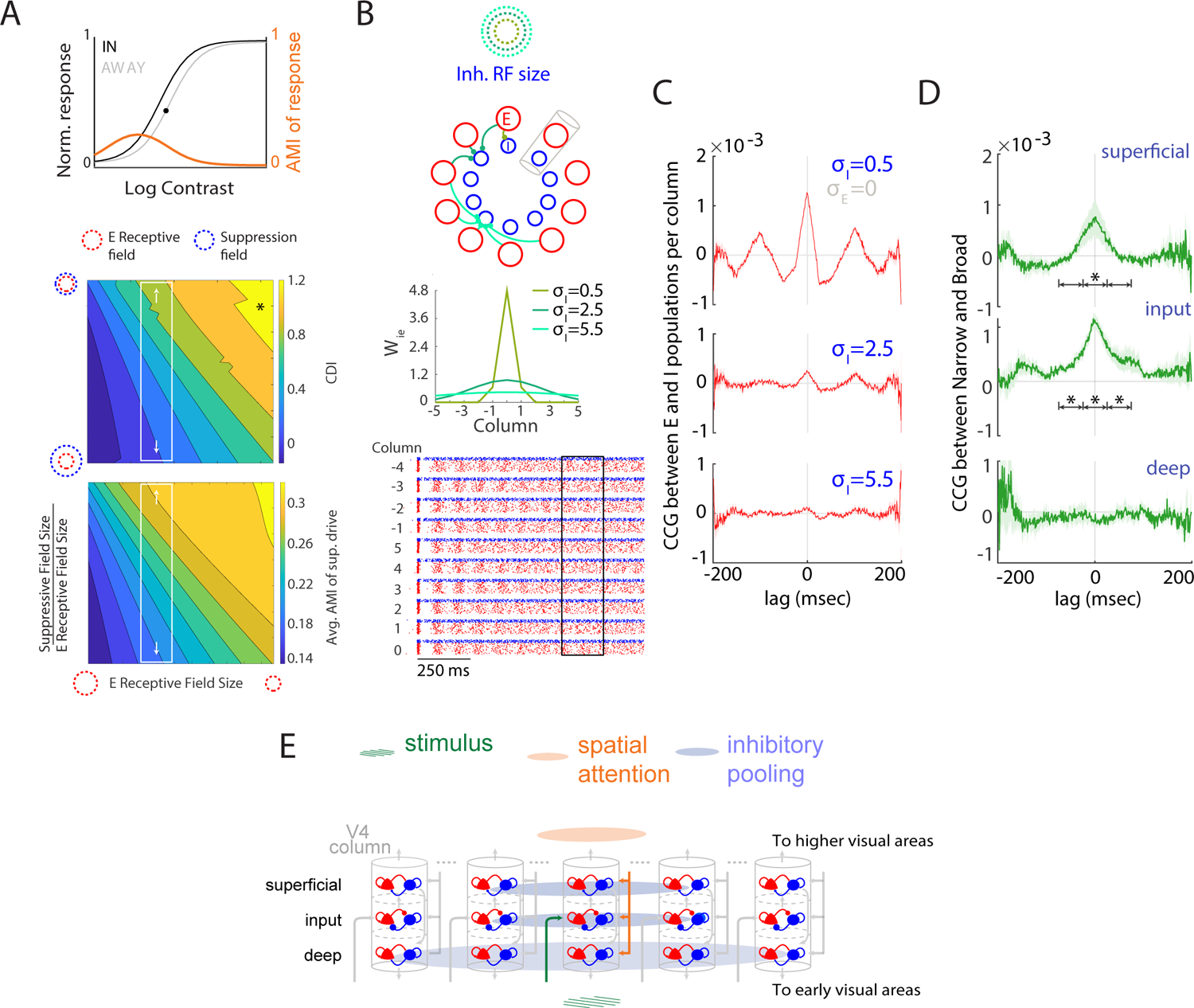
Computational Models Provide A Parsimonious Explanation for the Laminar Profile of AMI Contrast Dependence. (A) Predictions from the normalization model of attention with different suppressive field sizes or different excitatory (E) receptive field sizes. The top panel shows contrast response functions for a simulated neuron in the normalization model, when attending to a stimulus within the neuron’s receptive field (black curve) and when attending toward the opposite hemifield (grey curve). The orange curve represents the AMI. The black dot shows the inflection point of “attend away” responses that was used to delimit the low- and high-contrast ranges. The middle panel shows CDIs for simulated neurons as a function of the E receptive field size and the suppressive field size while holding the stimulus size and the attention field size fixed. The white rectangle depicts a potential mechanism that leads to the observed variation of CDIs across layers (change in suppressive field size). The black asterisk corresponds to the model parameters used for the simulation above. The bottom panel shows AMIs of suppressive drive as a function of the E receptive field size and the suppressive field size. For both simulations, the attention field size is 30 and the stimulus size is 5. The normalization model predicts the AMI of the suppressive drive to be correlated with the CDI of neuronal responses. (B) Simulations of a conductance-based E-I network with different inhibitory receptive field size. The schematic of the E-I networks corresponding to the possible mechanism in (A) are shown on top. 800 E and 200 I units were evenly distributed in 10 local E-I networks or “columns” around a ring. We interpret the normalization model’s suppressive field as the receptive field of inhibitory neurons in the E-I network model. E and I populations from the same column are mutually coupled. We modeled inhibitory receptive field size as the standard deviations (σ_I_) of E-I connections (W_ie_) across columns (middle panel, −5 and 5 are the same column). We changed the range of E-I connections across columns (W_ie_, shades of green) while keeping other within-column connections the same (including W_ee_, W_ii_, W_ei_). At the bottom, the raster plot shows the spiking activity for all units organized by their column IDs (blue, I; red, E) in response to a step input. The box depicts a 200 ms window used for computing cross-correlations between E and I populations. (C) Cross-correlograms between E and I populations in the same column with different inhibitory receptive field sizes. Cross-correlations were calculated using the pooled spike trains of E units and I units from the same column across 500 repeats of identical simulation and averaged across 10 columns. A larger inhibitory receptive field reduces the cross-correlation between local E and I populations. Mean ± SEM. (D) Cross-correlograms (mean ± SEM) between Narrow and Broad cell classes in the superficial, input, and deep layer. Cross-correlations were calculated using the pooled spike trains of Narrow class (putative inhibitory) neurons and Broad class (putative excitatory) neurons, and were averaged across sessions. The arrows mark 3 time intervals during which cross-correlations were averaged and compared between the superficial (or input) and the deep layers. Asterisk: The mean difference of cross-correlations between layers in the interval has a 95% CI above 0. For the estimation plot, see Supp. Figure 4B. (E) Proposed E-I networks in V4 accounting for the layer-wise CDI variations. The empirical data and the model simulations imply a larger inhibitory pooling size in the deep layer than those in the superficial and the input layers. The arrows depict the canonical information flow pathways in a columnar circuit.

The normalization model predicts the AMI of the suppressive drive (Figure 4A, bottom panel) to be correlated with the CDI of neuronal responses (Figure 4A, middle panel) (Reynolds and Heeger, 2009). However, the suppressive field in the model can be implemented by various biophysical mechanisms (Carandini, 2004). One possible mechanism is shunting inhibition via lateral connections from other neurons in the cortical neighborhood (Carandini and Heeger, 1994; Carandini et al., 1997; Kouh and Poggio, 2008), in which case the receptive field of local inhibitory neurons can approximate the suppressive field. Since the average AMI of the putative inhibitory (Narrow) cluster and CDI of putative excitatory (Broad) clusters in the input and deep layers in our empirical data (Figure 3D right panels, Figure 3E) is also correlated, we further explored this mechanism mediated by local inhibitory neurons. Under this assumption, the prediction about the changes in suppressive field size down the cortical depth from the normalization model transforms into one about changes in the excitatory (E) - inhibitory (I) connectivity along the cortical depth. Similarly, the prediction about the changes in excitatory receptive field sizes down the cortical depth can also transform into one about the changes in the E-E connectivity along the cortical depth (Gilbert and Wiesel, 1985; Hirsch and Gilbert, 1991). The layer-specificity of cortical connectivity implies different temporal signatures of neural activity across layers.

We next used a spiking network model to examine the effects of excitatory and inhibitory receptive field sizes on spike-time correlation between populations of local excitatory (E) and inhibitory neurons (I). Our spiking network model focuses on connectivity mechanisms for generating variable sizes of suppressive and excitatory receptive fields in a cortical network. The amplitude of the spike-time correlation between neurons has been shown to depend on both the connection strength and the background synaptic noise (Ostojic et al., 2009). Therefore, the spike-time correlation between neurons can be a proxy for the size of the postsynaptic neuron’s receptive field. We hypothesized that a smaller receptive field of the postsynaptic neuron would make the local connections more dominant against background inputs and lead to a higher spike-time correlation between the locally connected neurons. We examined how spike-time correlations change as a function of the inhibitory or excitatory receptive field size in a conductance-based model of spiking neurons (see Methods). We set up 10 local networks or “columns” of E and I units that were interconnected in a ring formation (Figure 4B, Supp. Figure 4C). Neurons within the same column were mutually coupled, while interactions between columns were confined to excitatory connections to local E and I neurons whose strengths decayed with distance between columns. All connections occurred with a probability of 0.5. We modeled the receptive field size as the standard deviation (σ_I_ or σ_E_) of the connection strength between columns (Figure 4B, Supp. Figure 4C). We performed simulations that generated spiking activity in response to a step input (Figure 4B, bottom panel). The spike-time correlation between local E and I populations was calculated using pooled spike trains within the same column; the resulting spike-time correlation was averaged across columns. We found that the inhibitory receptive field size has a critical impact on the spike-time correlation amplitude in such a network (Figure 4C), while the excitatory receptive field size has little effect (Supp. Figure 4C). A larger inhibitory receptive field (larger values of σ_I_) leads to a lower spike-time correlation between the local E and I populations in the network (Figure 4C). This result suggests that the prediction about inhibitory receptive field sizes down the cortical depth as the basis of CDI variation of broad-spiking neurons can be tested by examining the spike-time correlation between local E and I populations within each layer.

To test this prediction in our dataset, we computed the session-averaged spike-time correlation between Narrow (putative inhibitory neurons) and Broad (putative excitatory neurons) single units within each layer (see Methods). We found that the spike-time correlation amplitudes were higher in the superficial layer and the input layer than that in the deep layer (Figure 4D). We compared the spike-time correlations in the deep layer with those in either superficial or input layers, averaged within 3 different 50ms time windows. The 95% confidence interval of the mean difference between layers in either comparison was greater than 0 for the center window (Supp. Figure 4E). In accordance with our findings from the E-I network models (Figure 4C), this suggests that inhibitory neurons in the deep layer exhibit relatively broader receptive fields, which supports the prediction by the normalization model of attention (Figure 4A, middle panel). Our findings thus provide a parsimonious explanation for the layer- and cell-class specific contrast dependence of attentional modulation observed in area V4 (Figure 4E).

## DISCUSSION

Spatial attention plays a critical role in sensory guided behavior. It is thought to achieve this by enhancing the responses to low contrast stimuli in mid-tier visual cortical areas such as V4. While later stages of the visual processing hierarchy are thought to benefit from this manipulation, V4 also sends feedback projections to early visual areas that use veridical representation of contrast to aid object recognition. How area V4 meets these distinct information processing demands is not known. Contrary to the simplifying assumptions of prior empirical studies, we tested the hypothesis that V4 customizes its output to different stages of the visual processing hierarchy through layer- and cell-class specific attentional modulation of contrast computations. Recent advances in experimental techniques have shown layer- and cell-class specific functional specificity of computations in the cortical circuit (Adesnik and Naka, 2018; Adesnik and Scanziani, 2010; Naka and Adesnik, 2016; Olsen et al., 2012). However, these studies have been limited to species in which higher cognitive functions, such as attention, are challenging to study. Using computational approaches on laminar neural data in area V4 of the macaque, we find that the attentional modulation of neural responses to visual luminance contrast is indeed layer- and cell-class specific. We classified neurons into three functional cell classes defined by their action potential widths (Figure 2B); these classes show specificity in their response time to visual input (Figure 2D), attention effects on their contrast response functions (Figure 2F) and the contrast dependence of attentional modulation (Figure 3C). Specifically, narrow- and medium-spiking neurons show contrast-independent response modulation across layers; broad-spiking neurons, the putative projection neurons, exhibit significant contrast dependence of attentional modulation in the superficial layers, that project to higher level visual areas, but not in the deep layers, that project to earlier visual areas (Figure 3D, E). Notably, this highly significant laminar difference was not observable without cell-class identification (Supp. Figure 3C). These results provide the first evidence for our broad hypothesis that attentional modulation of contrast computations in the visual cortex is heterogeneous across those cell classes and layers that project to distinct stages of the visual processing hierarchy. The qualitative nature of the attention modulation of contrast in our data is not only distinct but suggests optimization for the computational demands of the target stages. Selective boosting of responses to low contrast stimuli is compartmentalized to the superficial output layers that project representations such as extended contours and object surfaces to higher areas (see Roe et al., 2012 for a review). Contrast-independent scaling of neural responses is confined to the deep output layers. Neurons in these layers project back to early visual areas that are reliant on faithful representation of luminance contrast for low-level feature extraction. We speculate that the contrast-independent attentive feedback provides a spatial boost signal to early visual areas that do not receive direct inputs from attention control centers such as the frontal eye fields (Ungerleider et al., 2008). This also aligns with the predictive coding model of object recognition, wherein V4 is a higher-level area in the object recognition hierarchy that generates predictions of lower-level activity, without corrupting the sensory landscape that is needed for error correction (Rao and Ballard, 1999).

When interpreted within the framework of the normalization model of attention (Figure 4A), the layer-specific attention modulation predicts differences in the spatial pooling of local inhibitory populations across layers. Such differences further predict a layer-specific signature of correlations between the activities of local inhibitory and putative excitatory neurons when explored in a spiking E-I network model (Figure 4B, C). We find robust evidence for differences in inhibitory spatial pooling across layers through our analyses of correlations between putative inhibitory and putative excitatory neurons in the superficial, input, and deep layers of the cortex (Figure 4D, E).

### Classification of cell-types

The duration of the extracellular spike waveform has been used to distinguish putative inhibitory interneurons from putative excitatory pyramidal cells in a wide range of species and across various brain regions (Ardid et al., 2015; Bruno and Simons, 2002; Constantinidis and Goldman-Rakic, 2002; Csicsvari et al., 1999; Fox and Ranck, 1981; Frank et al., 2001; Mitchell et al., 2007; Nandy et al., 2017; Rao et al., 1999; Simons, 1978; Swadlow, 2003; Wilson et al., 1994). In terms of attention effects, narrow-spiking neurons show stronger attention-dependent increases in absolute firing rates and firing reliability than broad-spiking cells (Mitchell et al., 2007). Unsupervised clustering algorithms are also effective in identifying subpopulations of neurons with distinct functional properties (Ardid et al., 2015; Gouwens et al., 2019; Hawken et al., 2020). It is important to note that the clusters we distinguished based on spike width may not correspond to neuronal classes differentiated based on morphology or protein expression patterns (Migliore and Shepherd, 2005; Tasic et al., 2018; Zeng and Sanes, 2017). Two possible correspondences exist between the narrow-spiking neurons and interneurons, and between the broad-spiking neurons and pyramidal cells (Connors and Gutnick, 1990; McCormick et al., 1985; Nowak et al., 2003). We find differences in both the firing rate (Supp. Figure 2A) and the attentional modulation of firing rates (Figure 3A, D) between clusters, suggesting their different functional roles in attention-mediated visual processing. Crucially, these distinct functional roles are reflected by the differences in contrast dependence of attentional modulation.

### Circuit roles for identified cell classes

The response time analysis (Figure 2D, E) in our study revealed potential roles for the three cell classes in the local circuit (Figure 5). Medium-spiking neurons have the shortest response time in all three layers, suggesting that they might be putative excitatory neurons receiving direct visual inputs in every laminar compartment. The broad-spiking class, which exhibits longer neuronal latencies, suggest a role in integrating lateral and feedback inputs, in addition to potentially acting as projection neurons due to their maximal attentional modulation overall, and in both the superficial and deep layers. Narrow-spiking cells, given their intermediate response time, might represent the typical inhibitory interneurons that are mainly driven by local excitatory inputs, and provide inhibitory inputs to local excitatory neurons.

**Figure 5.**
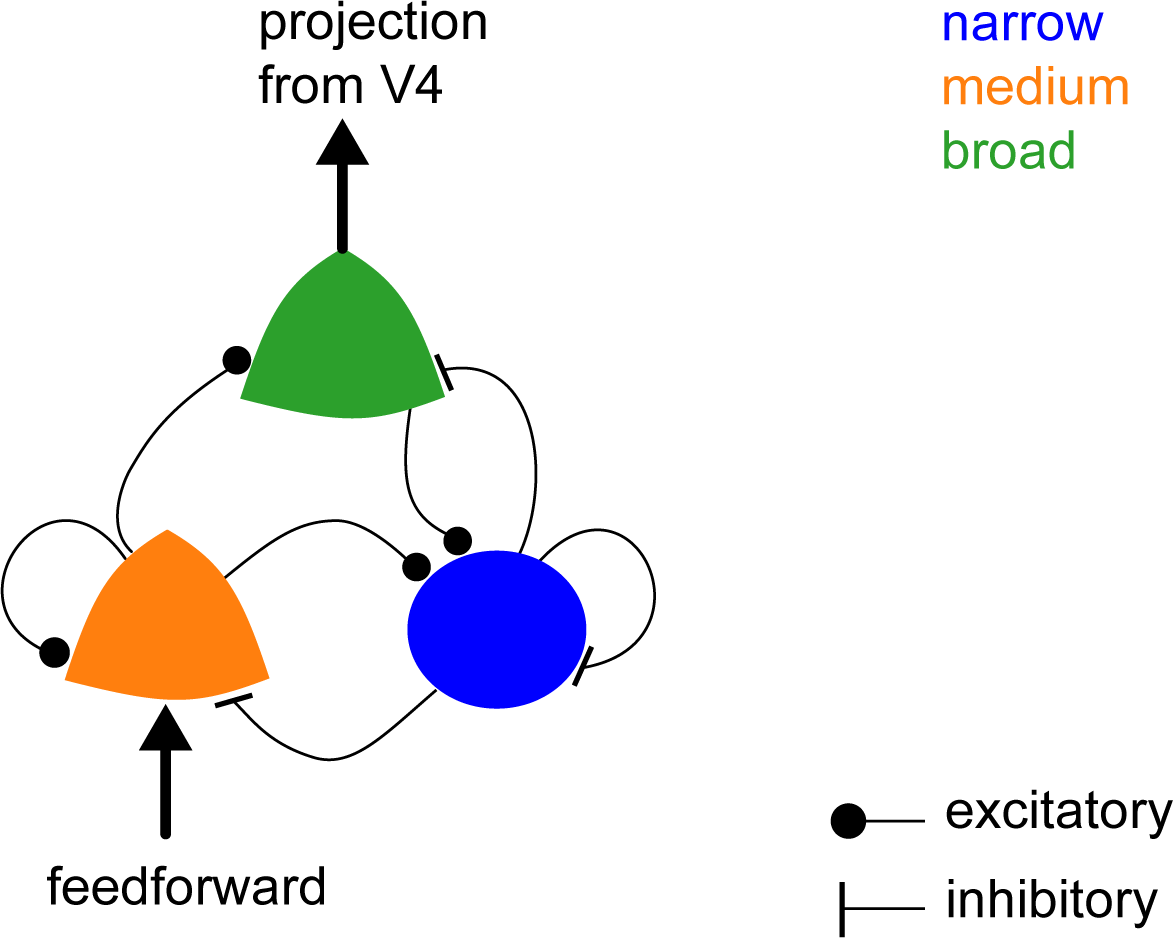
Predicted circuit roles for the narrow, medium and broad cell-classes in V4.

### Relation to prior studies of spatial attention in V4

Prior studies evaluating attention effects on neuronal contrast responses proposed either contrast-independent scaling of responses, termed as response gain (McAdams and Maunsell, 1999a; Morrone et al., 2002; Pestilli et al., 2009; Treue and Martinez Trujillo, 1999) or boosting of responses to low contrast stimuli, termed as contrast gain (Li and Basso, 2008; Li et al., 2008; Martinez-Trujillo and Treue, 2002; Reynolds et al., 2000) or an intermediate effect between the two (Huang and Dobkins, 2005; Williford and Maunsell, 2006). Although the overall attentional modulation of best-fitting CRF parameters in our dataset is consistent with the intermediate effect (Figure 1F, G), attention effects on individual clusters are variable: a mixture effect of response gain and contrast gain is observed for medium- and broad-spiking units (with differences in *c_50_* modulation); narrow-spiking cells show a response gain change (Figure 2F). Furthermore, broad-spiking neurons exhibit larger attention-dependent increases in response than the population mean, especially within the low-contrast range (Figure 3A). These observations suggest that attentional modulation of firing rate for broad-spiking cells may be more robust than that gleaned from previous studies that averaged across the whole recorded population. These cell-class specific increases in firing rate may significantly improve the signal-to-noise ratios of individual neurons, and therefore, act as another important contributor to the improvement of psychophysical performance due to attention in addition to reductions in correlations (Cohen and Maunsell, 2009; Mitchell et al., 2009).

### Our interpretation of the normalization model

The predictions from the normalization model (NM) of attention provide one possible explanation for the diverse contrast modulation patterns across layers. NM assumes both stimulus parameters and attention condition to contribute to the normalization input to local excitatory neurons. The stimuli presented in our experiments were optimized for the recording site and did not change with attention condition, and hence are not assumed to contribute differentially to the normalization mechanism. NM also assumes the sizes of attention field of the population to contribute to the normalization input to individual neurons. The attention field in NM describes the attention gain for each neuron in the population and depends on the animal’s attentional strategy employed during the experiment (Herrmann et al., 2010). The neural substrate for the attention field is unspecified in the NM, but we assumed the attention field to be constant across the cortical depth since the data was collected using a fixed experimental paradigm. However, given a lack of the biophysical mechanism underlying attentional modulation, our understanding of the attention field may be subject to future revision. The extent of excitatory receptive field, also termed as the stimulation field, in the NM can be mediated by various cortical connectivity patterns. While we explored a lateral pooling mechanism as the determinant of the receptive field extent of neurons, innervation specificity of feedforward synaptic input could be an alternative mechanism (Bruno and Simons, 2002; Hubel and Wiesel, 1962).

The variation in contrast dependence of attentional modulation observed across layers and cell classes (Figure 3D, E) in our data is explained by the NM in a most parsimonious way via the variability of the suppressive field size (Figure 4). However, the NM is agnostic to the neural machinery dedicated to the formation of neuronal tunings or the implementation of attentional modulation. To explore the implications of its field size predictions on spike-time correlations in a biophysical model, we considered the model’s stimulation field as the receptive field of putative excitatory projection neurons in a column, and its suppressive field as the receptive field of local inhibitory interneurons.

We implemented a spiking network model to relate the NM’s predictions of variable suppressive field sizes to variations in spike-time correlations in our data. It is important to note that our model is not a spiking network implementation of the entirety of attention computations described by the NM. The suppressive field in NM, which mediates divisive normalization, is a computation that can be can be implemented through a variety of mechanisms (see Reynolds and Heeger, 2009 for review). We chose one of the candidate suppression mechanisms – pooling of lateral inputs by local inhibitory interneurons (Carandini and Heeger, 1994; Carandini et al., 1997; Troyer et al., 1998). A feedforward mechanism of variable suppressive fields would yield a very similar prediction for spike-time correlations between local E and I populations. Our choice was guided by excellent agreement between the NM model AMI predictions and modulation patterns of related clusters in the input and deep layers. It is, however, important to note that in the superficial layers, putative inhibitory neurons (Narrow cluster) lack significant attention modulation in spite of robust boosting of responses to low contrast stimuli in putative excitatory neurons (Broad cluster). This does not agree with the predictions of the normalization model. There are three possible explanations for this observation: 1. Suppressive drive to broad-spiking neurons in superficial layer is not provided by local inhibitory neurons within that layer. 2. Superficial layer broad-spiking neurons inherit their contrast dependent attention modulation from the input layer. 3. Suppressive drive to broad-spiking neurons in the superficial layer is provided by non-PV local inhibitory neurons within the layer. Since PV neurons are a majority of the local interneuron population which itself occupies roughly 20% of the total neural population in the cortex, it is highly possible that our recordings did not sample the other inhibitory neuronal types. Indeed, studies from the mouse visual cortex suggest that SOM+ neurons play a key role in mediating lateral inhibition to layer 2/3 pyramidal neurons (Adesnik et al, 2012). Further studies are needed to distinguish the contributions of local vs feedforward computations to the attention effects in superficial layers.

When testing the model’s predictions in our dataset, we ascribed the stimulation field to the two non-Narrow clusters, including the Broad cluster identified in our layer-specific CDI analysis (Figure 3E). We ascribed the suppressive field to the receptive field of the Narrow cluster (putative interneurons).

## Conclusion

Attention increases the signal detection abilities of individual neurons. Whether the attention mediated firing rate variability is unchanged (McAdams and Maunsell, 1999b) or reduced (Mitchell et al., 2007), the response gain alone results in improved signal-to-noise ratio of individual neurons, and enhances the discriminability of the attended signal (McAdams and Maunsell, 1999b; Verghese, 2001). Attention mediated increases in neural responses to low- and intermediate-contrast stimuli can extend the separation between the neuron’s stimulus-evoked responses and its spontaneous activity, thereby improving the neuron’s sensitivity to low-contrast stimuli. There has, however, been a long-standing debate regarding the nature of interactions between attention and visual scene contrast that mediate object recognition. Previous theoretical studies have sought to resolve this based on the nature of differences in experimental paradigms (Reynolds and Heeger, 2009). Our work has exploited advanced experimental techniques to bring novel understanding of these interactions. Superficial cortical layers in area V4 that project to higher object recognition stages exhibit enhancement of low contrast stimuli. Deep layers that project to earlier visual areas exhibit contrast independent attentional scaling of neuronal responses. By identifying the compartmentalization of attention modulation among cortical layers, our study has uncovered a new dimension: the nature of interactions between attention and contrast is aligned with the demands of the visual processing hierarchy. A previous study has suggested that encoding of scene contrast and spatial attention by distinct neural populations in area V1 could fulfill its visual processing demands in the face of contrast dependent attentional feedback (Pooresmaeili et al., 2010). Our work has revealed an elegant mechanism of meeting these needs via laminar compartmentalization of attention modulation in area V4 that contributes to this feedback. Low-frequency synchrony between the thalamus and visual cortex has been suggested to guide the higher-frequency synchronization of inter-area activity that is critical to the communication of attention signals between brain areas (Saalmann et al., 2012). A contrast-independent effect of attention in the deep layer of V4 may also drive alpha rhythms of pulvino-cortical loops irrespective of stimulus conditions and maintain the transmission of attentional priorities across the cortex. Future studies are needed to test these and related hypotheses about the different functional roles of contrast-attention interactions in different cortical layers.

## METHODS

### Attention Task and Electrophysiological Recording

Well-isolated single units were recorded from area V4 of two rhesus macaques during an attention-demanding orientation change detection task (Figure 1A). The task design and the experimental procedures are described in detail in previous studies (Nandy et al., 2019; Nandy et al., 2017). While the monkey maintained fixation, two oriented Gabor stimuli were flashed on for 200 ms and off for variable intervals (randomly chosen between 200 and 400 ms). The contrast of each stimulus was randomly chosen from a uniform distribution of 6 contrasts (c = [10%, 18%, 26%, 34%, 42%, and 50%]). One of the stimuli was located at the receptive field overlap region of the recorded neurons and the other at an equally eccentric location across the vertical meridian. At the beginning of a block of trials, the monkey was spatially cued to covertly attend to one of the two spatial locations using instruction trials in which only one stimulus was presented. One of the two stimuli changed in orientation at an unpredictable time (minimum 1s, maximum 5s, mean 3s). The monkey was rewarded for making a saccade to the location of orientation change.

95% of the orientation changes occur at the cued location, and 5% occur at the uncued location (foil trials). We observed impaired performance and slower reaction times for the foil trials, suggesting that the monkey was indeed using the spatial cue to perform the task. The difficulty of the task was controlled by changing the degree of orientation change (randomly chosen from the following: 1°, 2°, 3°, 4°, 6°, 8°, 10°, and 12°). If no change occurred before 5 s, the monkey was rewarded for holding fixation (catch trial, 13% of trials).

While the monkey was performing the attention task, we used artificial dura chambers to facilitate the insertion of 16-channel linear array electrodes (“laminar probes”, Plexon, Plexon V-probe) or single tungsten microelectrodes (FHC Inc) into cortical sites near the center of the prelunate gyrus. Neuronal signals were recorded, filtered, and stored using the Multichannel Acquisition Processor system (Plexon). Neuronal signals were classified as either isolated single units or multiunit clusters by the Plexon Offline Sorter program. For the data collected from linear array electrodes, we used current source density analysis (Mitzdorf, 1985) to identify the superficial (Layers 1-3), input (Layer 4), and deep (Layers 5 and 6) layers of the cortex based on the second derivative of the flash-triggered LFPs (Bollimunta et al., 2008; Schroeder and Lakatos, 2009; Schroeder et al., 1998; Nandy et al., 2019; Nandy et al., 2017). Cell bodies of single units with bi-phasic action potential waveforms were assigned to the same layer in which the electrode channel was situated during recordings. Units that had tri-phasic waveforms or other shapes were excluded from analyses. Extracellular data were collected over 32 sessions (23 sessions in monkey A, 9 in monkey C) using linear array electrodes and 42 sessions (24 sessions in monkey A, 18 in monkey C) using single tungsten electrodes, yielding 410 single units in total (337 units using linear array electrodes and 73 units using single tungsten electrodes). Unit yield per session was considerably higher in monkey C than monkey A, resulting in a roughly equal contribution of both monkeys toward the population data.

### Contrast Response Function (CRF)

Neuronal responses were analyzed only for correctly performed trials, excluding instruction trials. We restricted all data analysis to non-target stimuli because neuronal responses to target stimuli were generally affected by the behavioral response or the reward delivery, which occurs on correct trials after the target’s appearance. Moreover, the larger number of non-target stimuli compared to target stimuli provided a more reliable response strength measure. For both attention conditions, the firing rate of a single unit in response to a particular contrast was measured by counting the number of spikes within a period of 60-260 ms after stimulus onset. Its baseline firing rate in each attention condition was extracted from a 200 ms window before a stimulus flash. The mean firing rates and the standard deviations (SDs) were generated across all stimulus flashes. We considered a neuron as visually responsive if any contrast responses exceeded its baseline firing rate by 4 SDs for both attention conditions. We found that 255 of 410 single units were significantly driven by the task stimuli.

We drew 1000 random samples of contrast responses from a normal distribution with the same mean and standard deviation as the experimental data for each visually responsive single unit. For each attention condition, we computed the CRF for each random sample by applying an ordinary least square fit to a hyperbolic ratio function:

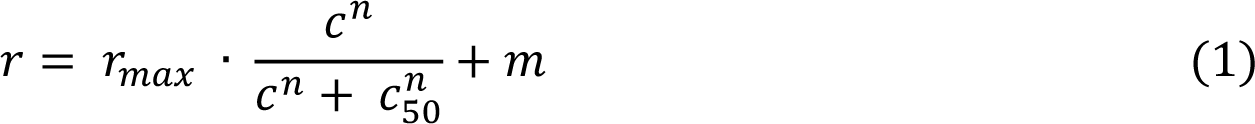

where *r* is the neuronal response, *r_max_* is the maximum attainable response, *c* is the contrast, *c_50_* is the contrast at which response is half-maximal, *m* is the baseline activity, and *n* describes the steepness of the response function and represents the neuron’s sensitivity to contrast. This function has been shown to provide a good fit to contrast response functions from visual cortices in cat and macaque monkey (Albrecht and Hamilton, 1982; Williford and Maunsell, 2006). We then averaged the best-fitting CRFs across random samples to generate the mean CRF for each visually responsive single unit (Figure 1C).

### Clustering Analysis

We used the *k*-means clustering algorithm (Hartigan and Wong, 1979) and a meta-clustering analysis (Ardid et al., 2015) to characterize cell classes based on peak-to-trough duration (PTD): we ran 500 realizations of the *k*-means for each *k* and selected the best replicate from 50 replicates for each realization. After 500 realizations of each *k*, we computed the probability that pairs of neurons belonged to a same cluster. Valid clusters were identified by setting a probability threshold (*p* ≥ 0.9). We considered clusters with at least five single units as reliable.

We computed the Akaike information criterion (AIC) and the Bayesian information criterion (BIC) to estimate the quality of clustering:

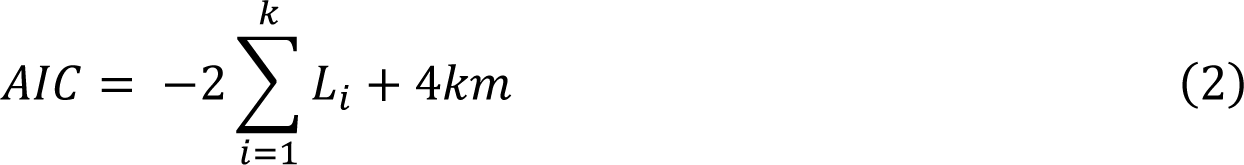

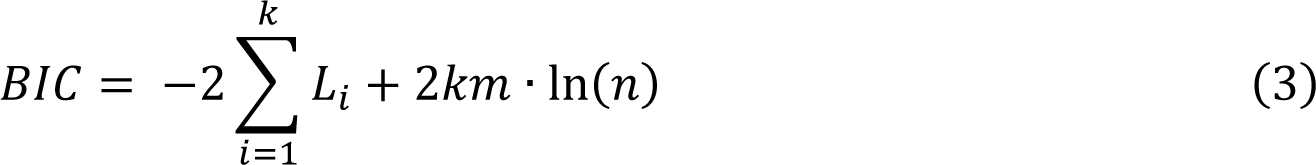

where *n* is the total number of units, *m* is the dimension of data in a being considered k-cluster solution (*m* = 1 in our study) and *L_i_* is the log-likelihood of sum of squares of cluster *i*.

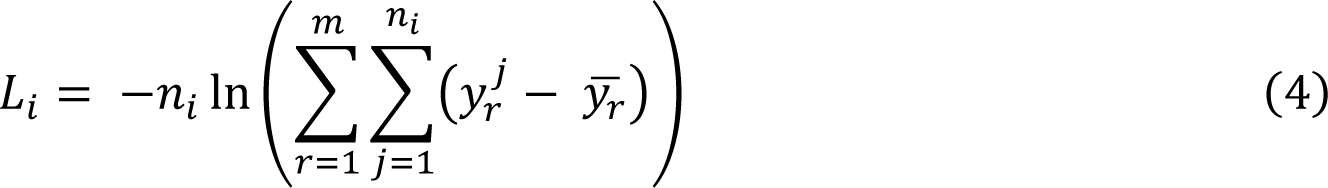

where *n_i_* is the number of units in cluster *i*, *y^j^_r_* is the *r^th^* feature value of unit *j*, y_r_ is the mean of the *r^th^* feature in cluster *i*.

To identify the most appropriate number of clusters, we implemented the Kneedle algorithm (Satopaa et al., 2011) to detect the point of maximum curvature (elbow point) in both AIC and BIC curves. The algorithm normalizes the points of the curve to the unit square and computes the set of differences between the *x*- and *y*-values, *i.e.*, the set of points (*x*, *y* − *x*) as in Figure 2A. An elbow point is then defined as the local maximum of the difference curve. Both AIC and BIC curves suggest that k = 3 is the optimal number of clusters for our dataset (Figure 2A).

### Firing Rate Estimation and Response Time

To measure a neuron’s response time to stimulus presentation, we pooled spikes across stimulus flashes for each single unit beginning 200 ms prior to stimulus onset and ending 260 ms after stimulus onset. We then convolved each spike with a Gaussian kernel and computed the average firing rates across flashes for each contrast level. The kernel bandwidth was variable at each time point selected by a bandwidth optimization algorithm that minimized the mean integrated squared error (Shimazaki and Shinomoto, 2010). Since attention barely affects neuronal latencies (Lee et al., 2007), we included trials from both “attend-in” and “attend-away” conditions to maximize the number of spikes. We only included units that fired at least 20 spikes within 200 ms before stimulus onset to ensure reliable firing rate estimates. The response time of a unit to the stimulus was defined as the delay after stimulus onset at which its average firing rate exceeded the maximal estimated firing rate (95% confidence interval of bootstrap samples) in the pre-stimulus period.

### Attentional Modulation Index and its Contrast Dependency

The attentional modulation index (AMI) of a neuron during the stimulus presentation with a specific contrast *c* was calculated using the best-fitting contrast response functions (*r*) from both attention conditions:

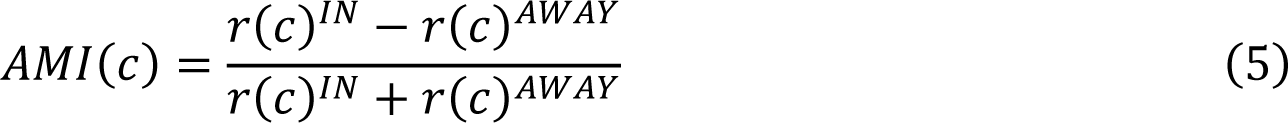

The contrast dependence of the AMI was measured by the contrast dependence index (CDI):

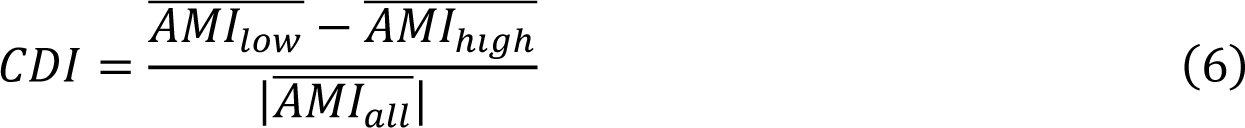

where 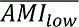 and 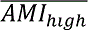 are the average AMIs within the low-contrast range and the high-contrast range, respectively. 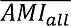 is the average AMI across all contrasts. *C_50_* from the best-fitting CRF during “attend away” condition delimited the range of low contrast (*c* < *c_50_*) and the range of high contrast (*c* ≥ *c_50_*). CDI measures how the AMI of a neuron fluctuates with the contrast of the stimulus. A zero CDI indicates that the AMI is independent of the contrast of the stimulus. More robust attentional modulation at the low-contrast range leads to positive CDIs, and more potent attention effects at the high-contrast range result in negative CDIs (Figure 3B). AMI and CDI were only calculated for those visually responsive neurons whose laminar locations were identified (n = 255).

### Normalization Model Simulations

We used the normalization model of attention (Reynolds and Heeger, 2009) to explore the neural mechanisms behind the variety of attentional modulation across layers (Supp. Figure 4A). The normalization model posits that the resulting firing rate (*R*) of the population of simulated neurons can be produced from a function of the stimulus drive (*E*), the attention field (*A*), and the suppressive drive (*S*):

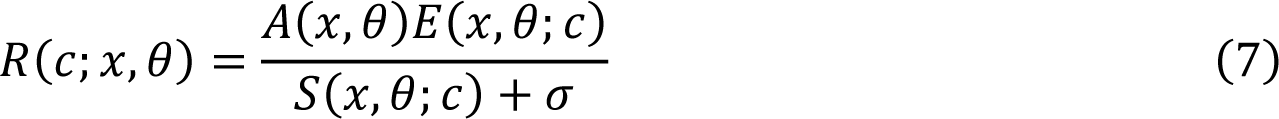

where *x* and θ represent the receptive field center and orientation preference of each neuron in the population. *c* is stimulus contrast and σσ is a constant that controls the contrast gain of the neurons’ response. The stimulus drive is derived from the stimulus and the stimulation field of the model neuron, which is its receptive field in the spatial and orientational space. The attention field describes the strength of attentional modulation as a function of receptive field center and orientation preference. The attentional modulation is 1 for unattended space and is greater than 1 for a small range of locations around the attended stimulus. We computed the suppressive drive by pooling the product of the stimulus drive and the attention field over spatial positions and orientations:

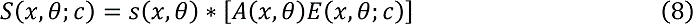

where *s*(*x*, θ) is the suppressive field and ∗ represents convolution. The stimulus, stimulation field (excitatory receptive field), attention field, and suppressive field all had Gaussian profile in space and orientation.

For simulations in Figure 4A, the stimulus size was 5 and the attention field size was 30. The CDI pattern holds for a range of stimulus sizes and attention field sizes (Supp. Figure 4B). The excitatory receptive field size and the suppressive field size were varied according to their ratios relative to the attention field size. For a given pair of stimulus size and attention field size, we changed the ratio of the attention field size to the excitatory receptive field size from 0.5 to 3 and the ratio of the suppressive field size to the excitatory receptive field size from 1 to 6. The orientation tuning width of the excitatory receptive field was 60^°^, and the suppressive field was nonspecific. A baseline activity of 0.5 was added after the normalization. For each combination of parameters, the AMIs were calculated using the model neuron responses from two attention conditions. The CDIs of AMIs were computed from the average AMIs within the low-contrast range and the high-contrast range delimited by the CRF’s inflection point from the “attend away” condition. For simulations in Supp. Figure 4B, we further modified the stimulus drive of the model to have either a nonlinear or an attention-modulated contrast response function. The nonlinear function was implemented as

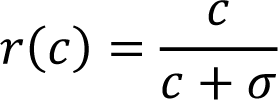

where σ is 0.26, matching the average *c_50_* of our data. We also applied either a multiplicative response gain (10% of increase in overall response) or a contrast gain (1% of increase in perceived contrast) to test the effects of different attention modulation of inputs on the model neurons’ responses.

### Computational Model

We set up a conductance-based model of *N*_E_ excitatory (E) and *N*_I_ inhibitory (I) neurons with a connection probability of 0.5 (Figure 4B). Neurons were evenly divided into 10 columns or local E-I sub-networks around a ring with the following within-column synaptic weights:

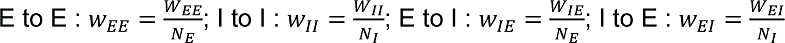

We only modeled E to I connections and E to E connections between different columns. The synaptic weights fell off with column distance following a Gaussian profile:

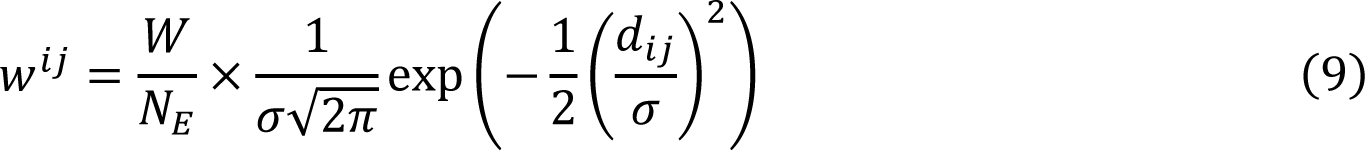

where w^ij^ is the synaptic weight between two columns (w^ij^_IE_ or w^ij^_IE_) and *d*_ij_ represents the distance from column *j* to column *i*. σ controls the pooling size of the postsynaptic inhibitory (σ_I_) or excitatory (σ_E_) neuron.

We simulated models of *N*_E_ = 800 excitatory and *N*_I_ = 200 inhibitory spiking units. The spiking units were modeled as Izhikevich neurons (Izhikevich, 2003) with the following dynamics:

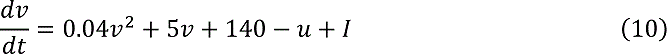

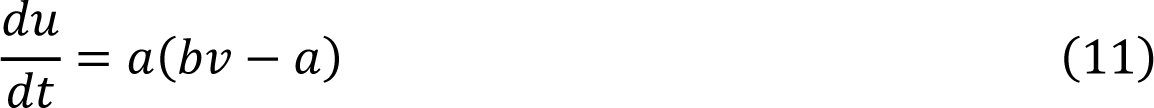

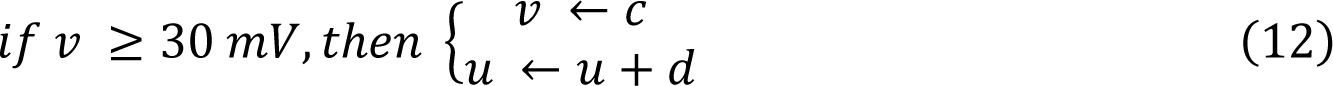

*v* represents the membrane potential of the neuron and *u* is a membrane recovery variable. *I* is the current input to the neuron (synaptic and injected DC currents). The parameters *a, b, c*, and *d* determine intrinsic firing patterns and were chosen as follows:

Regular spiking excitatory units: *a* = 0.02, *b* = 0.2, *c* = −65, *d* = 8

Fast spiking inhibitory units: *a* = 0.1, *b* = 0.2, *c* = −65, *d* = 2

Presynaptic excitatory neurons generate fast (AMPA) and slow (NMDA) synaptic currents, while presynaptic inhibitory neurons generate fast GABA currents:

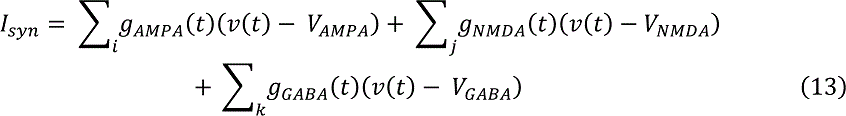

where *V_AMPA_* = 0, *V_NMDA_* = 0, *V_GABA_* = −70 are the respective reversal potentials (mV). The synaptic time course g(t) was modeled as a difference between exponentials:

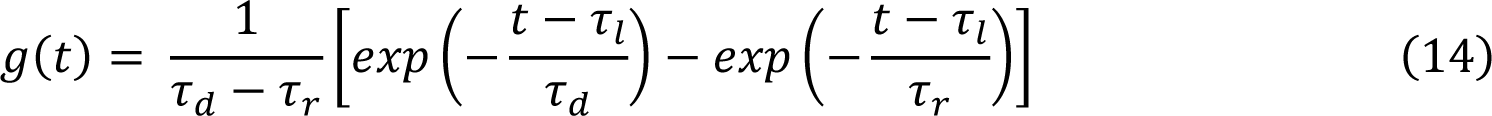

where the parameters τ_d_, τ_r_, and τ_l_ are the decay, rise, and latency time constants with the following values (Brunel and Wang, 2003): AMPA: τ_d_ = 2 ms, τ_r_ = 0.5 ms, τ_l_ = 1 ms; NMDA: τ_d_ = 80 ms, τ_r_= 2 ms, τ_l_ = 1 ms; GABA: τ_d_= 5 ms, τ_r_= 0.5 ms, τ_l_ = 1 ms; The AMPA to NMDA ratio is 0.45 (Myme et al., 2003).

We simulated the network with a DC step current (*I_DC_* = 4) of duration 1.2 s. Synaptic noise was sampled from a normal distribution (*I_syn-noise_*∼𝒩(μ = 0, σ = 3)). We pooled over spike trains of excitatory units and inhibitory units in each column separately and calculated the shuffled-corrected jittered cross-correlations from the two population spike trains binned at 1 ms within the 200 ms time window (800-1000 ms) after the initial transient response across 500 repeats of the simulation. Cross-correlations for different choices of σ_I_ or σ_E_ were reported as the average across columns (Figure 4C) (Harrison et al., 2007; Harrison and Geman, 2009).

### Spike Train Cross-correlations

The population cross-correlograms in Figure 4 report shuffled-corrected jittered cross-correlations (Harrison et al., 2007; Harrison and Geman, 2009). We computed the jittered cross-correlations by resampling two spike trains within a specific time window such that for each spike in the original data, a spike is chosen at random with replacement from within the same time window across trials, thus preserving the PSTH at the resolution of the jitter window. We computed the jittered cross-correlations with 4, 8, and 16 jitter windows, and the results of 8 jitter windows were shown. Shuffled cross-correlations were calculated by cross-correlating the first population spike train with the randomly permuted second population spike train. Both types of cross-correlations were averaged across trials and were further normalized by the geometric mean of the two spike trains’ firing rates and a triangular function that corrects for the amount of overlap for the different lags. The normalized shuffled cross-correlation was then subtracted from the normalized jittered cross-correlation to produce the shuffled-corrected jittered cross-correlation.

## AUTHOR CONTRIBUTIONS

MPJ & ASN conceptualized the project. XW analyzed the data, previously collected by ASN, and performed the computational modeling. MPJ supervised the project. XW, MPJ and ASN wrote the manuscript.

## ACKNOWLEDGEMENTS

This research was supported by NIH R00 EY025026 to MPJ, NARSAD Young Investigator Grant, Ziegler Foundation Grant and Yale Orthwein Scholar Funds to ASN, and by NEI core grant for vision research P30 EY026878 to Yale University.

## SUPPLEMENTARY MATERIAL

**Supp. Figure 1.**
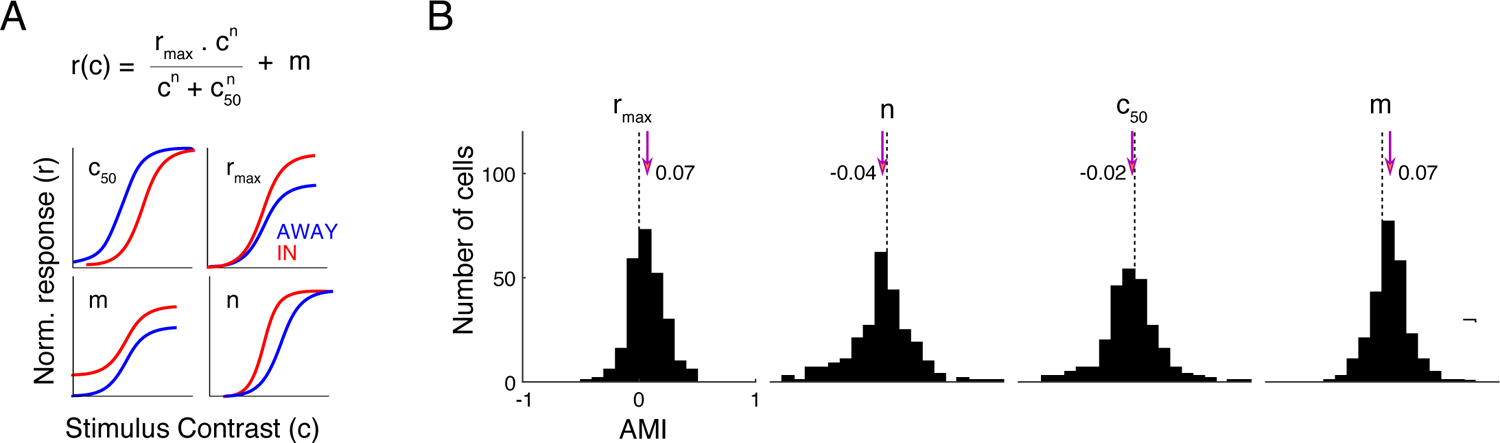
Attention Effects on Parameters from the Contrast Response Functions for All Neurons. (A) The mathematical function we used to fit neuronal contrast response functions is shown on the top. Schematics at the bottom show the effect of positive attentional modulation of each parameter on the contrast response functions. (B) Distributions of AMI of best-fitting parameters from the contrast response functions. The dashed lines mark the 0 modulation and the arrows indicate the median AMI values. The median AMI is significantly different from zero for each distribution (Mann-Whitney U test, p < 0.01).

**Supp. Figure 2.**
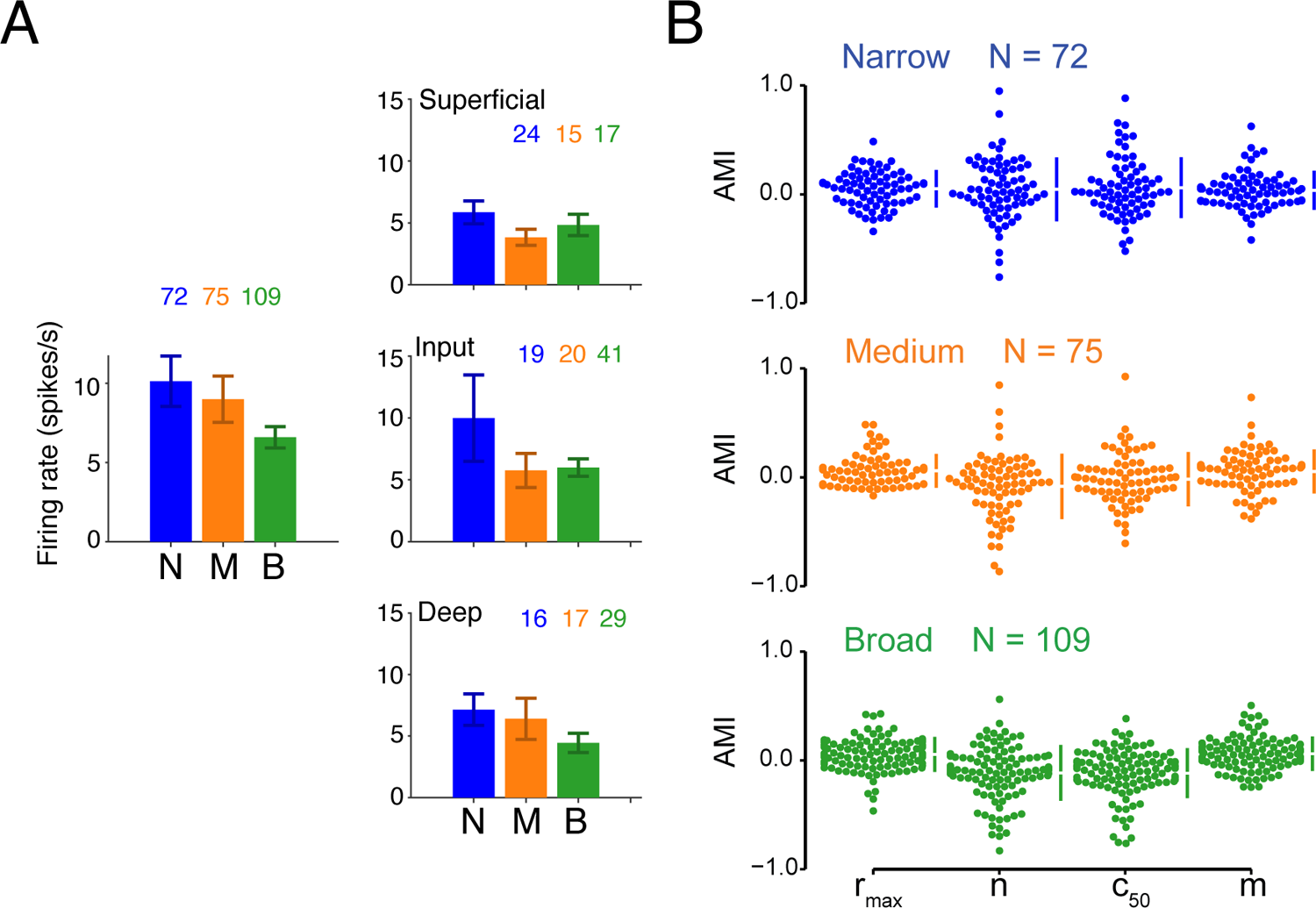
Cluster-wise Firing Rate and AMI of CRF Parameters. (A) Mean firing rate for visually responsive single units split by cell class or by layer. Neuronal firing rates were calculated from stimulus flashes with the highest common contrast across two monkey experiments in the “attend-away” condition. The number of single units within each cluster is shown. Mean ± SEM. (B) The raw data of AMIs of best-fitting CRF parameters for each cell class. The gapped lines to the right of each group show the mean and the standard deviations.

**Supp. Figure 3.**
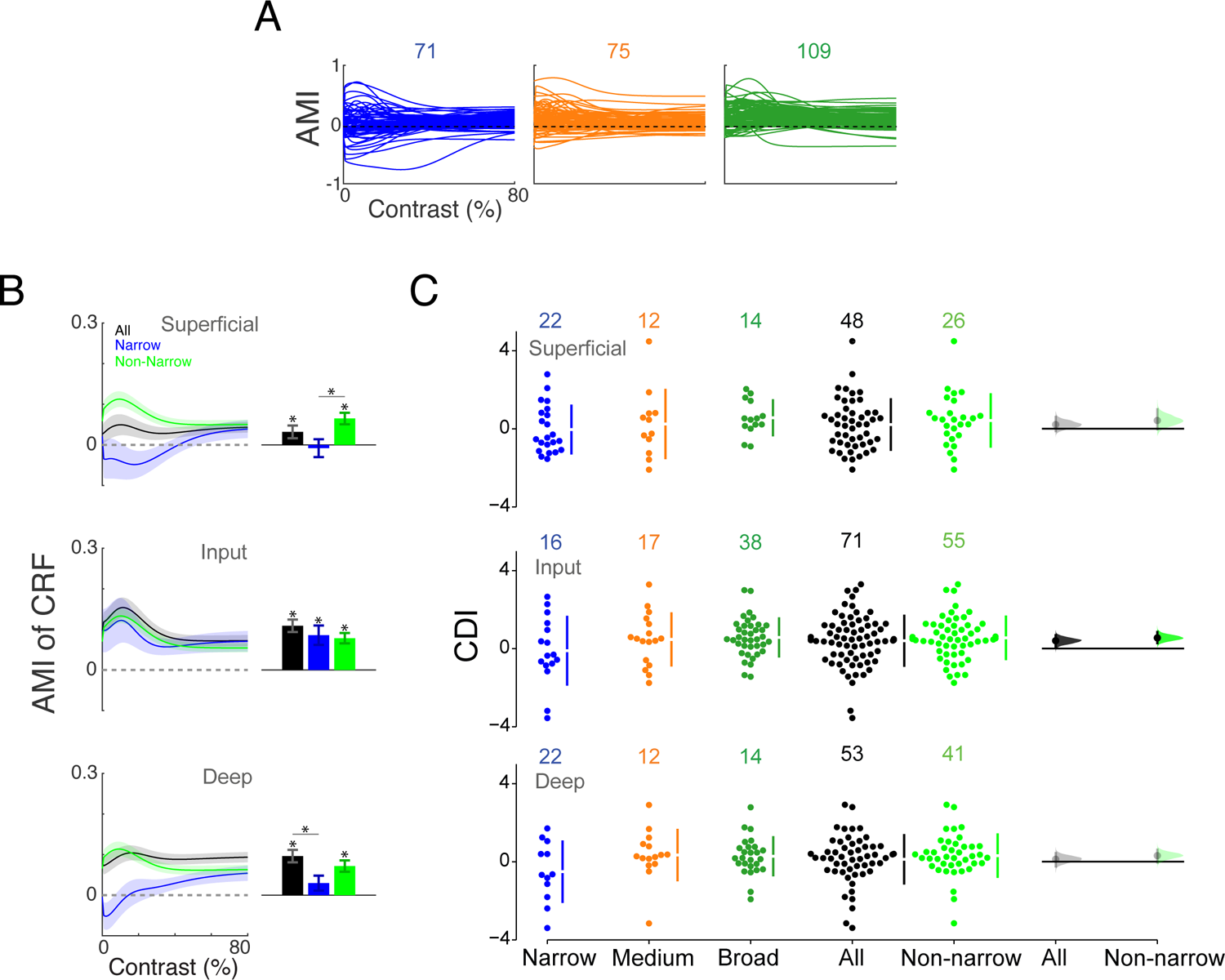
Raw Data of AMIs and CDIs for Each Cell Class. (A) The AMI as a function of contrast for individual units within each cell class. (B) Layer-wise AMI (mean ± SEM) for all units, Narrow unit, and non-narrow units as a function of contrast (left) or averaged across contrast (right). Asterisk indicates either the distribution is significantly different from zero or two distributions are significantly different (Mann-Whitney U test, p < 0.05). (C) The raw data of CDIs within each layer, including three clusters, the whole population and non-narrow units (Medium + Broad). The bootstrap distributions of mean CDI for the whole population and non-narrow units are shown on the right. Distributions with CIs inclusive of 0 are illustrated in faded colors.

**Supp. Figure 4.**
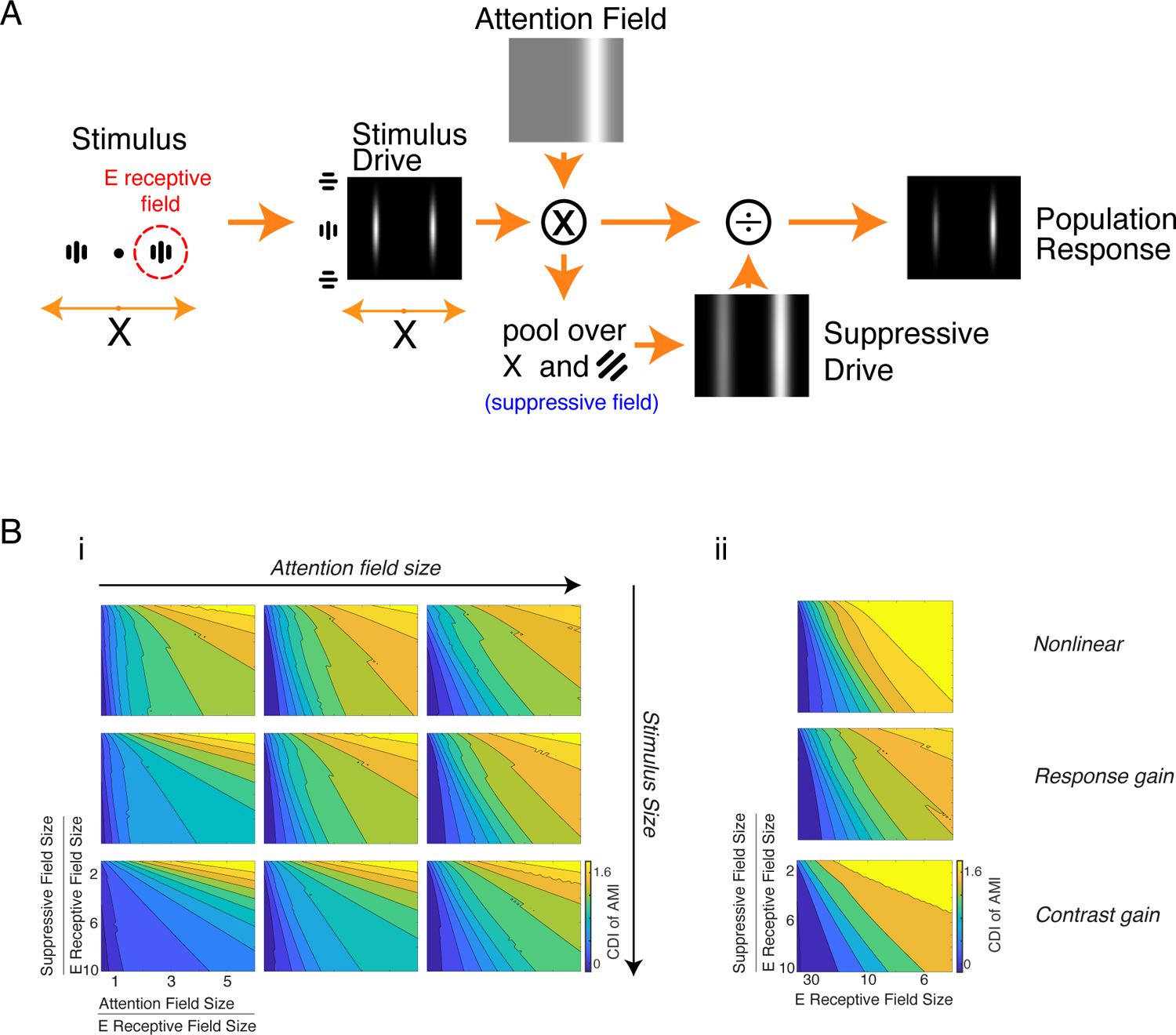
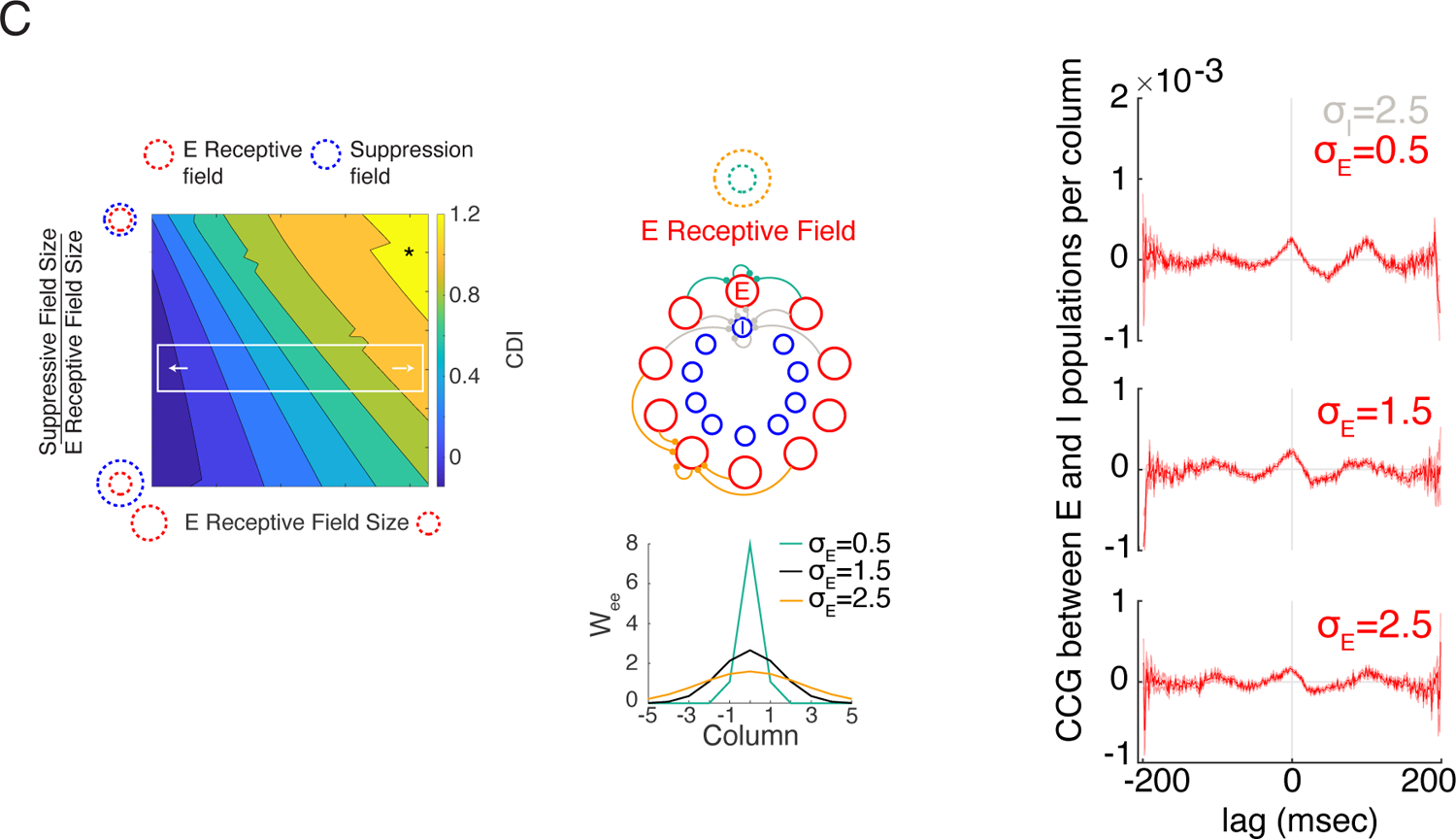
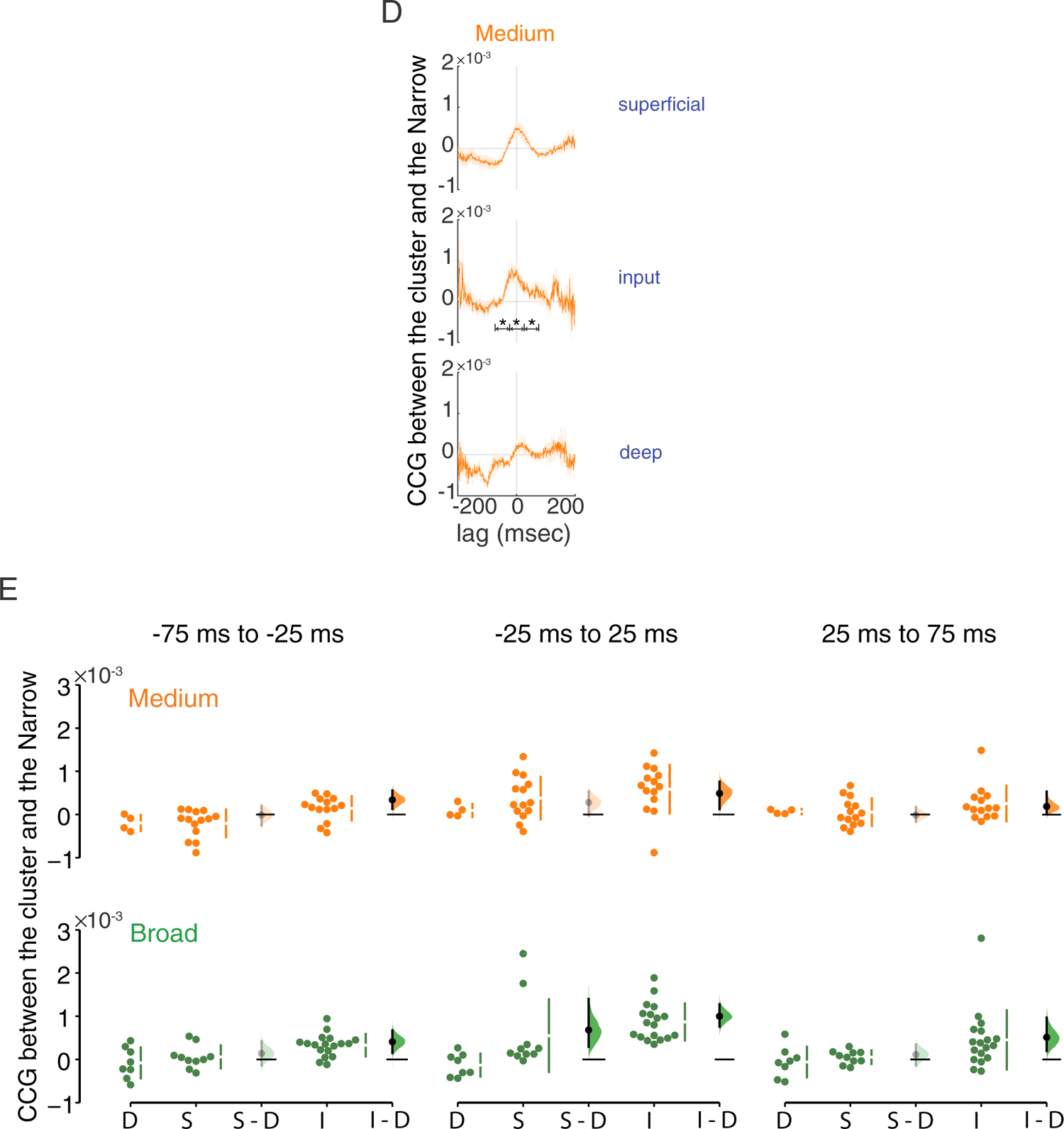
Normalization Model of Attention and CCG Analyses Between Cell Classes. (A) The structure of the normalization model of attention. The left panel shows a pair of orientated grating stimuli with identical contrasts, acting as input to the model. The central black dot indicates the fixation point. The dashed red circle indicates the receptive field of the model neuron centered on the grating stimulus. The stimulus drive shown in the middle panel is a collection of neural activity driven by the stimuli. Neurons are arranged based on their receptive field center (horizontal position) and orientation preference (vertical position). The values of the stimulus drive are shown by brightness. The top panel shows the attention field as a function of the receptive field center and the orientation preference. In this case, the attention is guided to the right stimulus position and does not vary with orientation. Grey areas indicate values of 1, and white areas indicate values greater than 1. The suppressive drive at the bottom is calculated from the point-by-point product of the stimulus drive and the attention field and then pooled over space and orientation according to the suppressive field size. The stimulus drive is multiplied by the attention field and then divided by the suppressive field to generate the output firing rates of model neurons (right panel). (B) i, CDIs for simulated neurons in the normalization model with different stimulus sizes and attention field sizes. In each panel, we vary the E receptive field size relative to the attention field size (x-axis), and the suppressive field size relative to the E receptive field size (y-axis). The pattern of CDI holds for a range of values of stimulus size and attention field size. ii, CDIs for simulated neurons in the normalization model with different types of inputs. We changed the stimulus drive input to the normalization model to have either a nonlinear or an attention-modulated contrast response function. We tested both the response gain (10% increase in overall response) and the contrast gain (1% of increase in detected contrast) effects. For these simulations, the attention field size is 30 and the stimulus size is 5. The pattern of CDI holds for different types of inputs. (C) Changes in E receptive field size (white box) can also lead to the variation of CDIs across layers (left panel). We tested this hypothesis in the E-I network by adjusting the standard deviation of between-column E-E connections (*W_ee_*) from narrow (green) to broad (orange) while keeping other connections the same (grey, including *W_ee_*, *W_ii_*, *W_ie_*) (middle panel). Cross-correlograms between E and I populations in the same column suggest that different E receptive field sizes have little impact on the spike-time correlations of local neural activity across layers (right panel). (D) Cross-correlograms (mean ± SEM) between Narrow and Medium classes in the superficial, input, and deep layers. Cross-correlations were calculated using the pooled spike trains of each cell class and were averaged across sessions. (E) The Cumming estimation plot shows the raw data of cross-correlation in each session and the mean difference of cross-correlation between the superficial (*S*) and deep (*D*) layers or between the input (*I*) and deep layers. We picked 3 time intervals (columns) to compute the cross-correlations of two pairs (Medium-Narrow, Broad-Narrow).

